# Ets-1 transcription factor regulates glial cell regeneration and function in planarians

**DOI:** 10.1101/2023.02.01.526519

**Authors:** Bidushi Chandra, Matthew G. Voas, Erin L. Davies, Rachel H. Roberts-Galbraith

## Abstract

Glia play multifaceted roles in nervous systems in response to injury. Depending on the species, extent of injury, and glial cell type in question, glia can help or hinder the regeneration of neurons. Studying glia in the context of successful regeneration could reveal key features of pro-regenerative glia that could be exploited for improvement of human therapies. Planarian flatworms completely regenerate their nervous systems after injury—including glia—and thus provide a strong model system with which to explore glia in the context of regeneration. Here, we report that planarian glia regenerate after neurons and that glia require neural structures to regenerate near the eyespot. We find that the planarian transcription factor-encoding gene *ets-1* promotes glial cell maintenance and regeneration. We also find that *ets-1*(RNAi) impairs nervous system architecture, neuronal gene expression, and animal behavior. Taken together, the discovery of *ets-1* as a regulator of glial persistence presents a critical first step in understanding glial regulation and potential roles of glia in planarian neurobiology. More importantly, we elucidate interrelationships between glia and neurons in the context of robust neural regeneration.

## Introduction

Glial cells are a heterogenous group of non-neuronal cells within nervous systems of many animals. Glial cells within the mammalian central nervous system (CNS) include astrocytes, oligodendrocytes, and microglia. Meanwhile, major glial types in the mammalian peripheral nervous system (PNS) include Schwann cells and satellite cells (Allen & Lyons, 2018; Jessen, 2004). Glial cells have been reported in most, but not all, bilaterian taxa studied (Hartline, 2011; Ortega & Olivares-Bañuelos, 2020). Molecularly, glial cells often express *glial fibrillary acidic protein* (GFAP), *glutamine synthetase* (GS), and/or *excitatory amino acid transporter* (EAAT), but these markers are not universal across species (Hartline, 2011). Functionally, glial cells play important and diverse roles in the development, maintenance, and activity of the nervous system across bilaterians. These roles include regulating neural cell numbers and migration; assisting in axon guidance; maintaining ionic homeostasis and neurotransmitter uptake; facilitating synapse architecture; and remodeling neural circuitry (Allen & Lyons, 2018; Jäkel & Dimou, 2017; Jessen, 2004; Oikonomou & Shaham, 2011; Shaham, 2015; Yildirim *et al*., 2019).

In regeneration, glial cells play dynamic roles that depend on the species, as well as the region, extent, and duration of the injury. For instance, mammalian astrocytes and microglia respond to injury through “reactive gliosis,” which can lead to formation of a “glial scar” near the lesion (Adams & Gallo, 2018; Anderson *et al*., 2016; Burda & Sofroniew, 2014; Escartin *et al*., 2021; Gallo & Deneen, 2014; Pekny *et al*., 2014). Glial scarring can promote neuronal survival, but can also limit axonal regeneration (Anderson *et al*., 2016; Myer, 2006; Rolls *et al*., 2009; Silver & Miller, 2004). In contrast, glial scarring does not occur in fish and insects due to the presence of bridging glia in the zebrafish spinal cord (Goldshmit *et al*., 2012; Mokalled *et al*., 2016), and phagocytic ensheathing glia in the adult *Drosophila* neuropil (Doherty *et al*., 2009; Purice *et al*., 2017). Glial roles in the context of successful neural regeneration are only beginning to be uncovered, partly due to the limited regenerative capacity of the CNS in many traditional model organisms (Alesci *et al*., 2022; Silver *et al*., 2015; Tanaka & Ferretti, 2009).

Freshwater flatworms called planarians undergo whole-body regeneration without scarring, including *de novo* regrowth and rewiring of the entire brain. Genes that include *intermediate filament* (IF-1), *calamari,* and *estrella* mark glia present in the planarian nervous system, opening an opportunity to explore glial biology in an organism with complete regenerative capacity for the first time (Roberts-Galbraith *et al*., 2016; Wang *et al*., 2016). Glia in the planarian CNS and PNS express genes that encode proteins for neurotransmitter uptake and metabolism (*e.g. solute carrier 1a- 5*/*EAAT*; *glutamine synthetase-1*), indicating an overlap in function with astrocytes in the mammalian CNS (Roberts-Galbraith *et al*., 2016; Wang *et al*., 2016). Previous work also determined that transcription factor *forkhead box protein factor-1* (Scimone *et al*., 2018) and Hedgehog signaling from ventral-medial neurons impact glial gene expression in planarians (Currie *et al*., 2016; Wang *et al*., 2016), though the consequences of changes in glial gene expression are not known. Furthermore, single-cell sequencing transcriptomic atlases suggest that planarian glia share similarities to other mesenchymal cell types that express *cathepsin*, including pigment cells, parenchymal cells, and other uncharacterized cell types (Fincher *et al*., 2018; Plass *et al*., 2018). Transcriptional similarity between *cathepsin^+^* cell types has been interpreted as a lineage-based relationship, though this has not yet been definitively shown. Many fundamental aspects of planarian glial biology remain unexplored, including how glia regenerate and what roles, if any, they play in the planarian nervous system during homeostasis and regeneration.

ETS transcription factor genes regulate gliogenesis in both vertebrates and invertebrates (Kiyota *et al*., 2007; Klämbt, 1993; Kola *et al*., 1993). *Drosophila pointed*, which encodes an ETS transcription factor, is necessary and sufficient for longitudinal and midline glial differentiation (Klaes *et al*., 1994). In *Xenopus*, *Ets-1* directly regulates radial glial formation and is important for neuron-glia interaction during embryogenesis (Kiyota *et al*., 2007; Klämbt, 1993). Human and mouse *Ets-1* are expressed in astrocytes of the cortex and are involved in astrocyte differentiation (Amouyel *et al*., 1988; Fleischman *et al*., 1995). In planarians, previous studies indicate that *ets-1* plays roles in several cell types, including pigment cells, glial cells, and other uncharacterized cathepsin^+^ cells (Dubey *et al*., 2022; He *et al*., 2017), but a definitive function for *ets-1* in glia has not yet been clearly established.

In this study, we determined that the transcription factor Ets-1 promotes maintenance of existing glial cells in uninjured tissues and regeneration of new glial cells after injury in the planarian *Schmidtea mediterranea*. Furthermore, using *ets- 1*(RNAi) to perturb glia, we investigated potential roles for planarian glia for the first time. We determined that *ets-1*(RNAi) non-cell-autonomously impacts neuronal gene expression, neuropil size, and animal movement. Taken together, our work demonstrates that planarian *ets-1* plays a conserved and crucial role in glial cells during regeneration and tissue homeostasis in planarians. Finally, this work explores spatiotemporal and functional relationships between planarian glial cells and neurons during regeneration and embryogenesis.

## Results

### Planarian glial cells arise after neurons

In vertebrate and *Drosophila* development, neurogenesis precedes gliogenesis (Anthony *et al*., 2004; Barnabé-Heider *et al*., 2005; Bayraktar & Doe, 2013; Klämbt & Goodman, 1991; Viktorin *et al*., 2011). We reasoned that understanding the sequence of neuronal and glial development and regeneration in planarians could help form testable hypotheses about glial cell specification and function. We first amputated planarians pre-pharyngeally and fixed animals at several time points to establish a timeline of neuronal and glial regeneration. Using a marker of cholinergic neurons, *choline acetyltransferase* (*ChAT*) (Nishimura et al., 2010), we saw re-establishment of neurons within a primordial brain around 3 days post-amputation (dpa) and clear brain organization at 5 dpa (Fig 1A, top; S1A). Our results were consistent with previous reports that new neurons are born beginning at 2 dpa (Cebrià *et al*., 2002; Inoue *et al*., 2004). The expression pattern of *ChAT* in the brain remained comparable from 5 dpa onward.

**Figure 1.**
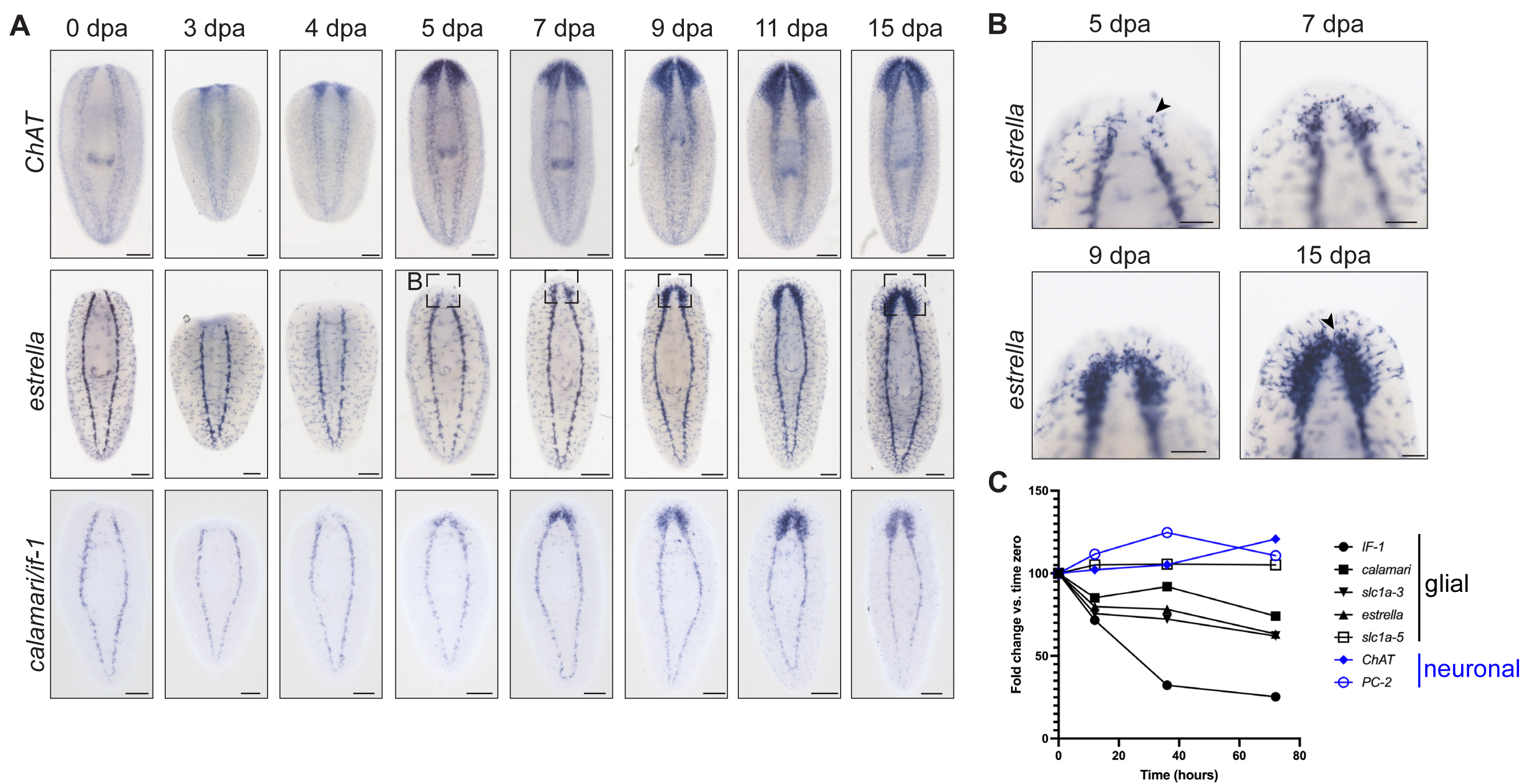
Planarian glial cells regenerate after neurons. (A) ISH regeneration timeline of neurons (top), marked by *choline acetyltransferase* (*ChAT*), and glial cells (bottom), marked by *estrella* and pooled *if-1*/*calamari*. Dashed boxes indicate insets shown in B. (B) Inset of *estrella* expression in head blastema at 5-, 7-, 9-, and 15 dpa. Arrowheads show round *estrella*^+^ cells that progress to stellate morphology. (C) RNA-seq of tail fragments regenerating head tissue illustrates glial and neuronal marker transcript levels at early time points post-amputation (Roberts- Galbraith *et al*., 2016). Planarian glial markers are downregulated in the first 72 hours post amputation. Ventral view, anterior up. Scale bars: 200 µm; insets 100 µm.

In contrast to *ChAT*, the earliest appearance of *estrella^+^* glial cells in the newly regenerated head occurred between 4-5 dpa (Fig 1A, middle) (Roberts-Galbraith *et al*., 2016). *estrella*^+^glia initially re-appeared in small numbers and were round; cell number increased over time and cells also adopted stellate morphology (Fig 1B). By 15 dpa, the distribution of *estrella*^+^ cells appeared identical to that in uninjured planarians. We confirmed this timeline for glial cell regeneration using two additional glial markers: *intermediate filament-1* (*if-1*) (Roberts-Galbraith *et al*., 2016; Wang *et al*., 2016) and *calamari* (Wang *et al*., 2016)(Fig 1A, bottom; S1A). *IF-1* is also downregulated after injury in RNA-sequencing (RNA-seq) data and we confirmed that other glial markers were transiently downregulated at early time points after injury (Fig 1C) (Roberts-Galbraith *et al*., 2016). Taken together, our results show that the planarian glial cells respond to injury by changing gene expression and regenerate in new tissue after neurons.

Next, we asked whether the temporal order of cell birth holds true in embryogenesis. Development of the adult nervous system begins during S5, with the expression of transcription factor-encoding genes with roles in neuronal subtype specification (Davies *et al*., 2017). Genes required for differentiated neuron function, including terminal selector genes required for neuronal subtype maintenance and genes involved in synapsis and neurotransmission, show enriched expression during Stage 6 (S6), Stage 7 (S7), and Stage 8 (S8) (Davies *et al*., 2017). Expression of the neuronal marker *pc-2* was detected by single embryo bulk RNA-seq as early as Stage 2 (S2) (Fig 2A). We did not detect robust expression of glial markers *if-1*, *calamari*, and *estrella* by bulk RNA-Seq prior to S7 (Fig 2A; S1B-C). One exception was *EAAT*/*slc1a-5*, which was first expressed during S6; however, single cell transcriptomic data from adult asexual planarians suggest that *EAAT* is also expressed in muscle and other *cathepsin*^+^ cells (Fincher *et al*., 2018; Plass *et al*., 2018).

**Figure 2:**
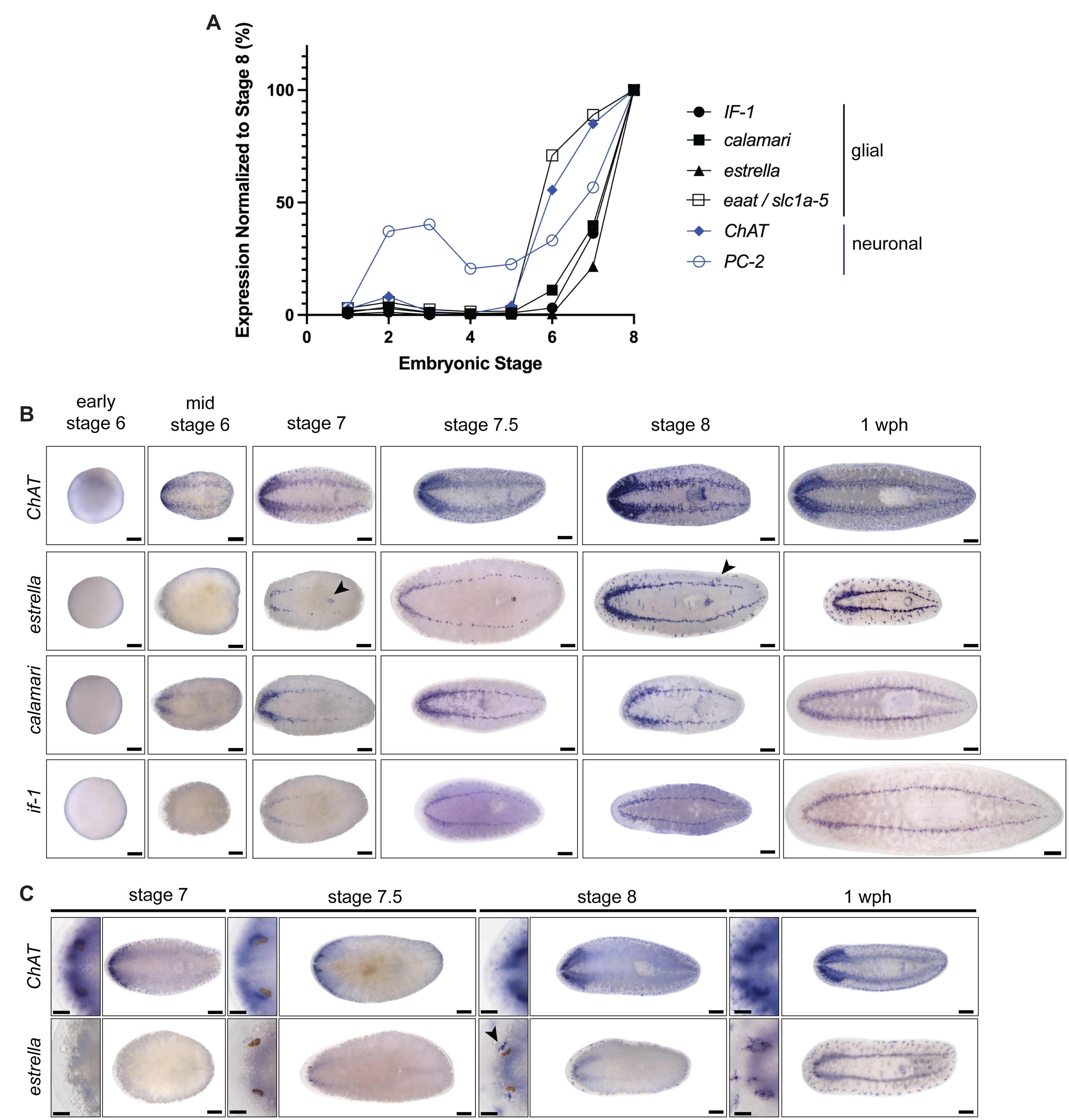
Planarian glial cells develop after neurons during embryogenesis. (A) Single embryo RNA-sequencing shows expression of neuronal and glial markers (Davies *et al*., 2017). Neuronal genes are enriched as early as S2 and expressed strongly at S6; glial gene expression initiates around S6 and strengthens at S7-S8. (B) ISH of neuronal (*ChAT*) and glial markers (*if-1*, *calamari*, *estrella*) in planarian embryos (early S6, mid-S6, S7, S7.5, S8) and juveniles (1 week post-hatching), ventral views. Arrowheads show *estrella* expression in mouth and peripheral nervous system. (C) ISH of *ChAT* and *estrella* on staged embryos, dorsal views. Insets show expression near and within the eyespot. *estrella* expression near the eye does not appear until S8 (indicated by arrowhead); *ChAT* expression, in contrast, is seen as early as S7.5 beneath the eyes. Anterior left. Scale bars: 200 µm; eyes 50 µm.

ISH was performed to examine spatiotemporal expression patterns for neuronal- and glial-enriched transcripts on staged planarian embryos and juveniles (1 week post-hatching; wph)]. By ISH, *ChAT*^+^ neurons can be seen at S6 (Fig 2B) (Davies *et al*., 2017). Neuronal markers *synaptotagmin* (*syt1-1*) and *pc-2* also appeared during S6 (Fig S1B-C). By the end of S7, neuronal markers highlighted well-developed brain and ventral nerve cord (VNC) structures that were nearly contiguous and extended to the posterior end of the embryo. Peripheral neurons were evident in both *ChAT* and *syt1-1* ISH during stage S6 and photoreceptor neurons were evident by S7 (Davies *et al*., 2017)(Fig. 2C, S1B-C). In planarians, as in other organisms, neurogenesis initiated in the anterior with the formation of the brain primordia, followed by VNC formation, which again showed early anterior bias (Fig 2B; S1B).

We next examined expression of glial markers during embryonic development. Markers specific for differentiated glial cells were expressed beginning in S6-S7 (Davies *et al*., 2017). Expression of *calamari* in the CNS initiated first and was detected by WISH in the brain primordia and anterior domain of the developing VNC at mid-S6 (Fig 2). *if-1* and *estrella* showed expression in the brain primordia and developing VNC in the anterior half of the embryo during S7 and in the posterior by S7.5 (Fig 2B). *estrella* was expressed near the mouth beginning at S7, around the eyes, and in putative PNS glial cells in S8 hatchlings (Fig 2B-C). All PNS glia and those around sensory structures like the eye appeared later than neurons. Expression of peripheral glial marker genes was heavily biased towards the ventral side of the embryos at all stages assayed (Fig 2B-C, S1C). Taken together, we determined that gliogenesis occurs after neurogenesis during embryogenesis and adult regeneration, like the relative order of cell birth in other animal species.

### Glial regeneration in sensory structures might depend on neurons

In many organisms, the interactions between neurons and glia during development are dynamic and reciprocal. It has been proposed that neurons can act as a “blueprint” for glial cell development by regulating the migration, survival, and proliferation of glial cells (Allen & Lyons, 2018). The birth order we established for planarian glia and neurons led us to seek ways to test whether glia depend on neurons for birth or final location. *estrella*^+^ glial cells are present in vicinity of the photoreceptor neurons of the eyespots and the ciliated sensory neurons along the dorsal midline and lateral margins of the body (Fig 3A, 3C, S2). We chose to examine neural dependence in these locations due to low, countable density of glia and the ability to perturb subsets of neurons in these locations.

**Figure 3.**
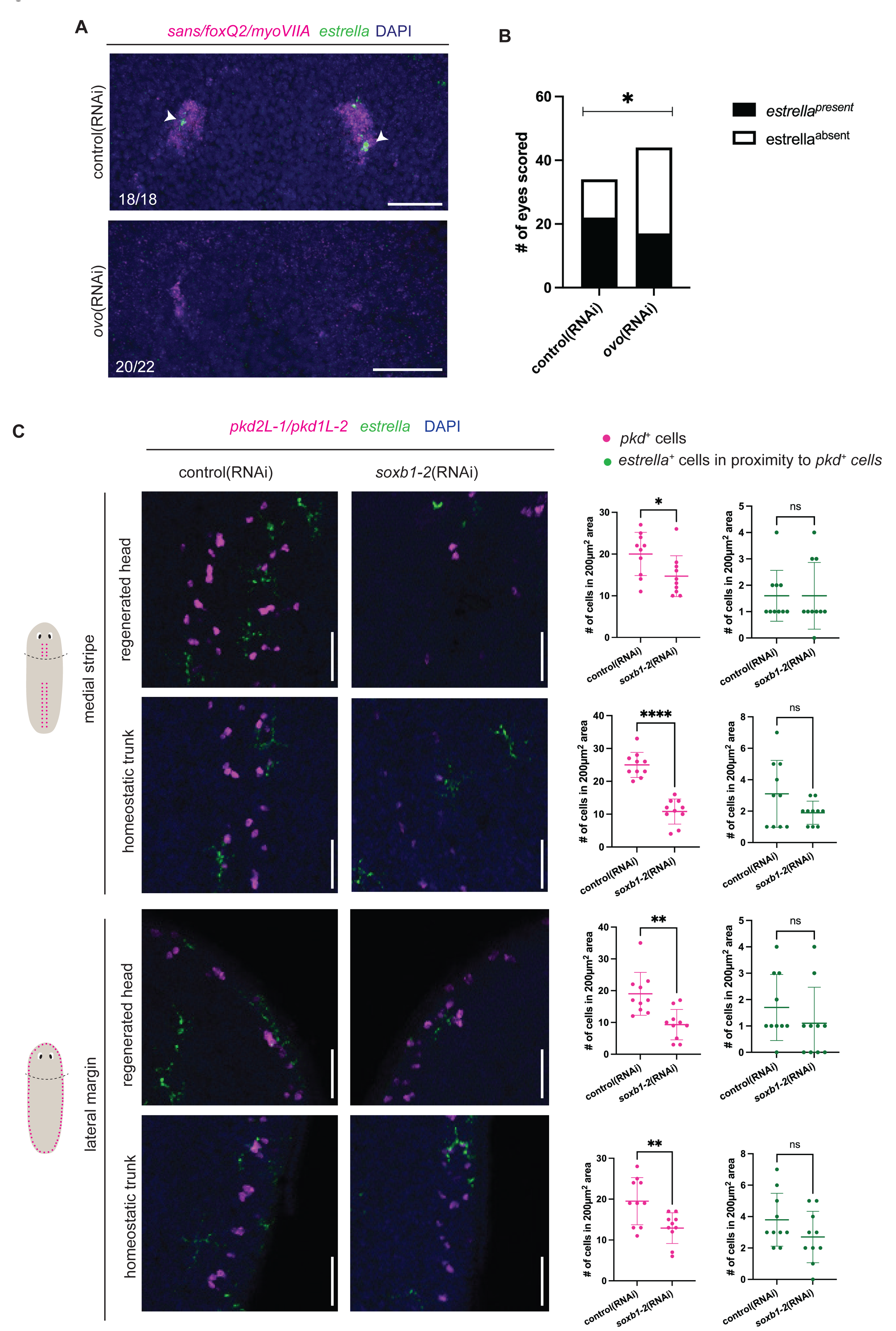
Presence of *estrella*^+^ cells near the eye is eye-dependent. (A) Regenerated control and *ovo*(RNAi) animals subjected to FISH with riboprobes detecting pooled *sans/foxQ2/myoVIIA* (photoreceptor neurons, magenta) and *estrella* (glia, green) and stained with DAPI (blue). Arrowheads indicate *estrella*^+^ cells in proximity of the eye. (B) Quantification of estrella^present^ (black) or estrella^absent^ (white) eyespots after control or *ovo*(RNAi). *p≤0.05 (Fisher’s exact test). (C) Control and *soxB1-2*(RNAi) animals subjected to FISH detecting pooled *pkd2L-1*/*pkd1L-2* (sensory neurons, magenta) and *estrella* (green), and nuclei (DAPI, blue) in medial stripe and lateral margins in regenerated and non-regenerated tissue. Quantification of *pkd*^+^ or *estrella*^+^ cells in respective regions to the right. *p-value≤0.05, **p-value≤0.01, ****p- value≤0.0001, ns=not significant (unpaired t-tests with Welch’s correction). n=10. Dorsal views. Scale bar: 50 µm.

To investigate whether *estrella*^+^ cells are dependent upon photoreceptors or pigment cells in the eye, we knocked down *ovo* to reduce eyespot regeneration (Lapan & Reddien, 2012). As previously reported, *ovo*(RNAi) caused reduced expression of pooled photoreceptor neuron markers *sans*/f*oxQ2*/*myoVIIA* (Fig 3A; S2A). In the same animals, we examined *estrella* expression in the typical location of the eyespots (Fig 3A-B) (Roberts-Galbraith *et al*., 2016). Compared to control, we saw significant reduction of *estrella*^+^ cells present in typical area of eyespots after *ovo*(RNAi) (Fig 3B). Our data suggest that the presence of photoreceptor neurons or pigment cup cells is required for the expression of regenerated *estrella*^+^ glial cells nearby.

We also sought to determine if glial regeneration and maintenance are dependent on sensory neurons present along the medial and lateral dorsal surfaces. Transcription factor *soxb1-2* is required for regeneration of many sensory neurons that function in these regions (Ross *et al*., 2018). We thus performed head amputations on *soxB1-2*(RNAi) animals (Ross *et al*., 2018), and examined *estrella* expression. As previously shown, pooled *pkd2L-1/1L-2*^+^ neurons decreased in number after *soxB1- 2*(RNAi) in the medial stripes and the lateral margin of the body (Fig 3C; S2B). In both newly regenerated and non-regenerating tissue, we did observe decreases in *estrella*^+^ cells near *pkd*^+^ sensory neurons in medial and lateral regions, however the differences were not significant (Fig 3C; S2B). However, we do acknowledge that perduring neurons present in these locations may still promote glial localization.

Taken together, our data show that reduction in cells of the eyespot leads to reduction in *estrella^+^* cells that localize in the region of the eye. Similarly, we saw a consistent but nonsignificant decrease in regeneration of PNS glia after perturbation of neurons in this location. These results suggest that population of the nervous system with glia during regeneration may be promoted by antecedent neuronal cell types.

### Putative transcription factor *ets-1* affects glial gene expression

Beyond the impact of neurons on glial regeneration, we also wished to identify genes required cell-autonomously for glial regeneration. Work in several other organisms established roles for ETS-family transcription factors in glial cell identity (Amouyel *et al*., 1988; Fleischman *et al*., 1995; Kiyota *et al*., 2007; Klaes *et al*., 1994; Klämbt, 1993). We utilized phylogenetic analysis to show that *S. mediterranea* Ets-1 is similar to *D. melanogaster* and *C. salei* Pointed (Chen *et al*., 1992; Klämbt, 1993; Pribyl *et al*., 1988) and mammalian and *Xenopus* Ets-1 (Fig S3) (Slupsky *et al*., 1998; Stiegler *et al*., 1990; Watson *et al*., 1988).

Planarian *ets-1* is expressed widely throughout the mesenchyme (Fig 4A) and has been reported to play roles in specification and maintenance of pigment cells and other *cathepsin^+^* cell types (Dubey *et al*., 2022; He *et al*., 2017). Prior studies on planarian *ets-1* reported either no role in impacting *estrella* expression (He *et al*., 2017) or decreased *calamari* expression in the head only (Dubey *et al*., 2022). To uncover the full impact of *ets-1* on glial cells, we optimized an RNAi paradigm by adjusting numbers and dosage of dsRNA feedings (Fig. S4A). Following optimization, we knocked down *ets-1* and examined glia with multiple markers (Fig 4B-C). *ets-1*(RNAi) animals showed a reduction of *estrella* expression during homeostasis and at 7 dpa (Fig 4D). We observed that the *estrella* expression pattern differed in three distinct ways in *ets- 1*(RNAi) animals. In both homeostatic and regeneration tissue, we observed gaps in *estrella^+^*signal in the VNC (Fig. 4D, black arrowhead; 4E); reduction of peripheral *estrella^+^*cell number (Fig 4D, red arrowhead; 4F-I); and decreased *estrella^+^*cell number in the newly regenerated head (Fig. 4D, bottom; S4B-C). Taken together, our data suggest that *ets-1* promotes maintenance of *estrella*^+^cells in pre-existing tissue as well as regeneration of new *estrella*^+^cells.

**Figure 4.**
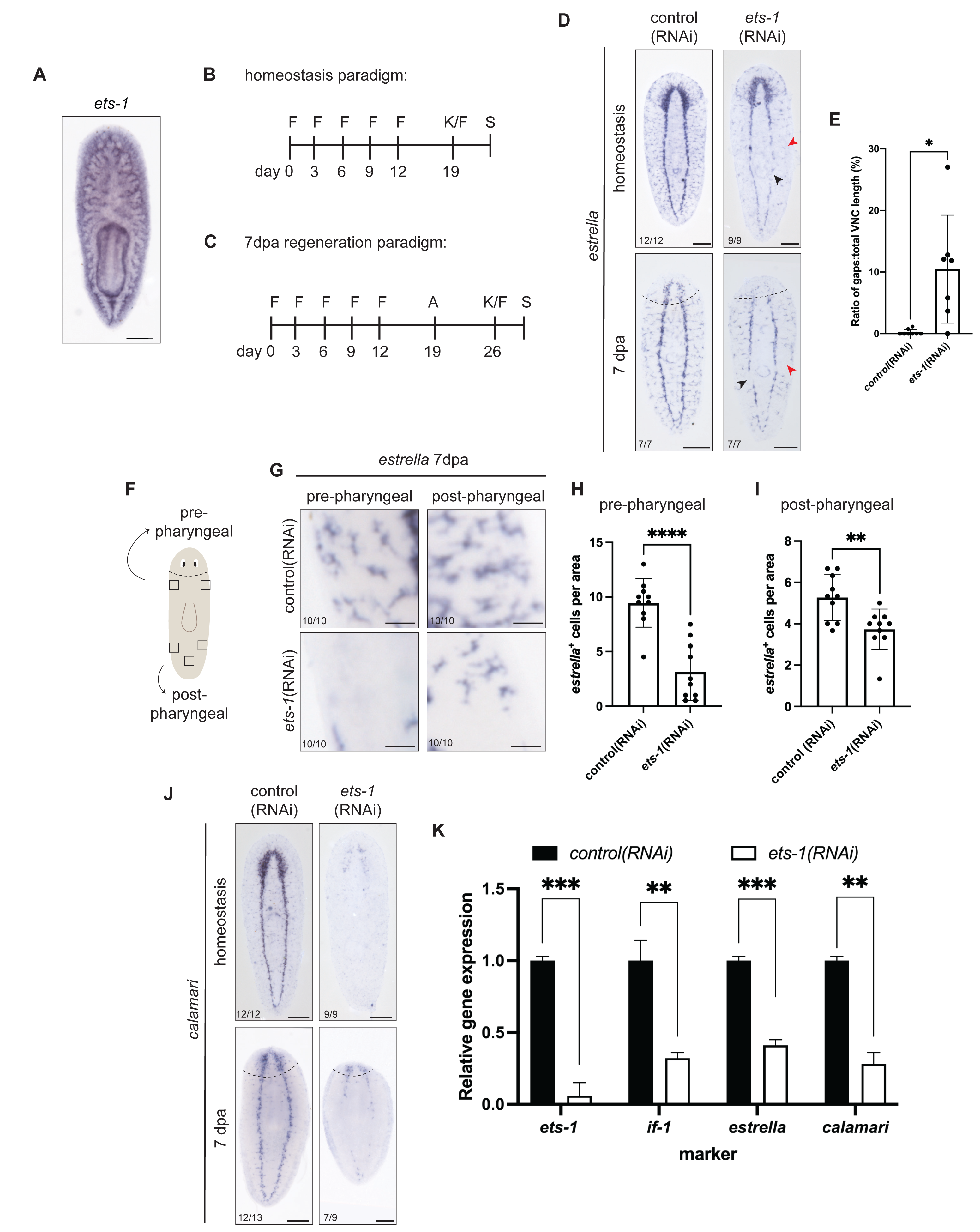
*ets-1* affects glial gene expression. (A) WISH of *ets-1* in uninjured, untreated animals. (B-C) RNAi feeding paradigm for *ets- 1*(RNAi) for homeostasis and 7 dpa regeneration. Feeding (F), amputation (A), kill/fix (K/F) and staining (S) are indicated. (D) ISH of *ets-1*(RNAi) animals detecting *estrella* expression. (E) Regenerated *ets-1*(RNAi) animals have higher stretches of VNC gaps compared to control (see also black arrowhead in D). *p≤0.05 (unpaired t-tests with Welch’s correction). (F) Illustration showing 200 µm^2^boxes drawn to quantify PNS *estrella*^+^ cell number. (G-I) Insets and quantification showing reduced peripheral *estrella^+^* cells in control and *ets-1*(RNAi) animals (see also red arrowhead in D). **p- value≤0.01,**** p-value≤0.0001 (unpaired t-tests with Welch’s correction). (J) ISH of *ets-1*(RNAi) animals detecting *calamari* expression. (K) RT-qPCR used to detect levels of *ets-1, if-1, estrella,* and *calamari* transcripts after RNAi. Error bars: SEM. **p- value≤0.01, ***p-value≤0.001 (unpaired t-tests). Dashed lines indicate amputation site. Ventral views, anterior up. Scale bar: (A, D, J) 200 µm, (G) 50 µm.

We next asked how *ets-1* impacts other glial markers. In comparison to control, *ets-1*(RNAi) worms exhibited reduced *calamari* expression throughout the body, in pre-existing and new tissue (Fig. 4J)(Wang *et al*., 2016). Interestingly, the reduction of *calamari* expression was strongest in the VNC and more dramatic than the reduction of *estrella*. We further quantified relative expression of *if-1*, *estrella*, and *calamari* transcripts in control and *ets-1*(RNAi) animals by RT-qPCR and observed significant reduction in all glial transcripts (59-72% reduction; Fig 4K).

Our previous data showed that changes in *estrella* and *calamari* expression after *ets-1*(RNAi) were not identical. We further investigated cell-specific effects of *ets- 1*(RNAi) using fluorescent *in situ* hybridization (FISH) (Fig 5A-C). As with ISH, we saw significantly reduced cell number using each glial marker (Fig 5B-C). Looking at the overlap in gene expression, we saw that there were significantly fewer *estrella*^+^/*calamari*^+^ and *estrella*^-^/*calamari*^+^ cells in regenerating heads (Fig 5D, F). However, there was not a significant difference in *estrella*^+^/*calamari*^-^ cells (Fig 5E). Our data suggest that in addition to impacting glial numbers overall, Ets-1 also influences gene expression in remaining glia, potentially reflecting additional roles in cell state or maturation. Overall, our data demonstrate a requirement for *ets-1* in glial cell maintenance, regeneration, and gene expression in planarians.

**Figure 5.**
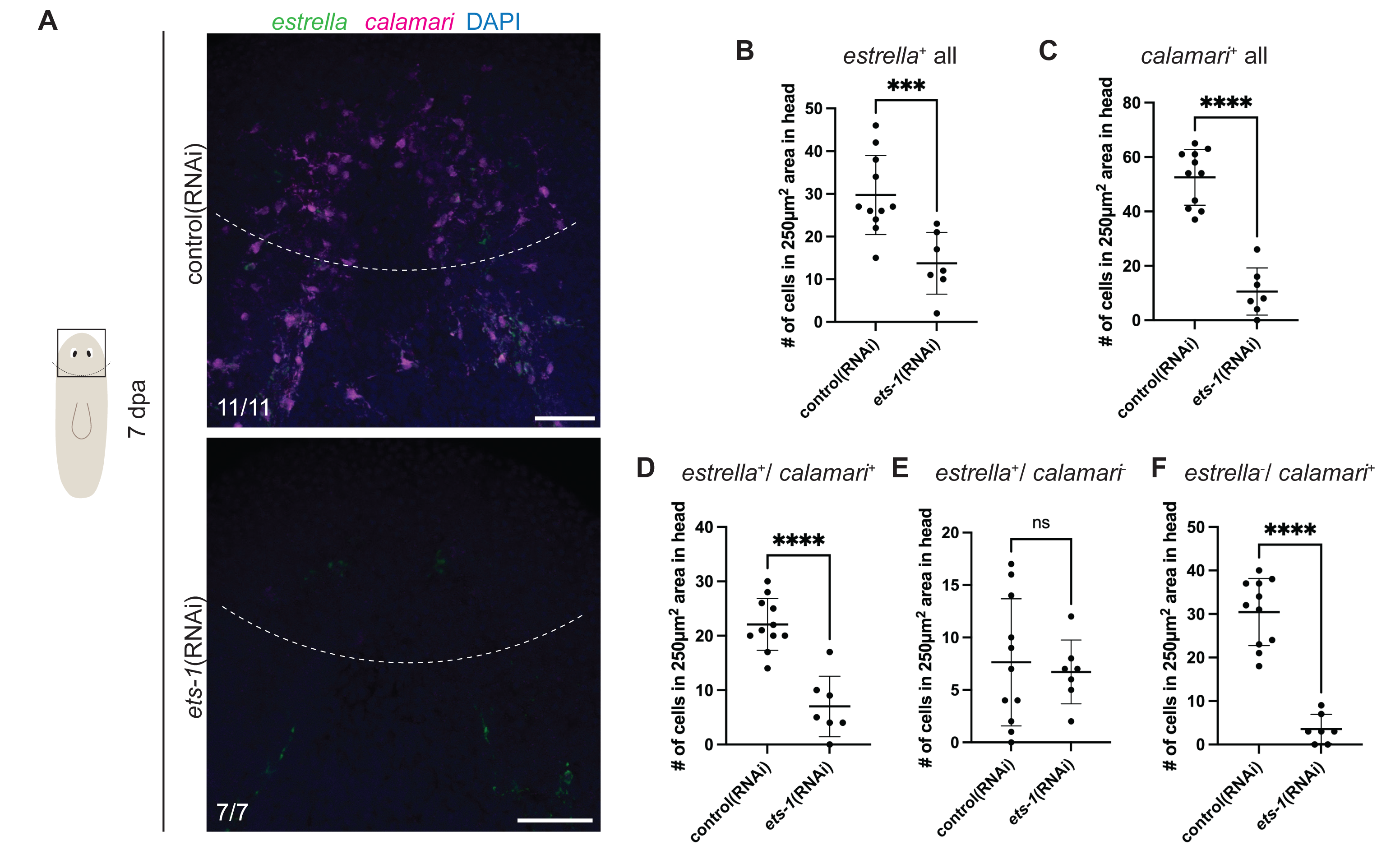
*ets-1* affects *estrella* and *calamari* differently. (A) FISH for *calamari* (magenta) and *estrella* (green) expression, with DAPI (cell nuclei, blue) in newly regenerated heads in control and *ets-1*(RNAi) animals. Dashed line indicates amputation site. (B-F) Quantification of cell expression patterns in 250 µm^2^ areas in head blastema in control and *ets-1*(RNAi) samples. ***p-value≤0.001, ****p≤0.0001, ns=not significant (unpaired t-tests with Welch’s correction). Scale bar: 200 µm.

### *ets-1* affects multiple *cathepsin*^+^ subclusters

Continuous knockdown of *ets-1* eventually leads to animal death (Fig S5)(Dubey *et al*., 2022). This led us to ask how *ets-1* affects cell types beyond glia and pigment cells (Dubey *et al*., 2022; He *et al*., 2017). Single cell RNA sequencing (scRNA-seq) atlases cluster planarian pigment cells and glial cells with other cells that express *cathepsin*(Fincher *et al*., 2018; Plass *et al*., 2018)(Fig S6A). We first confirmed previous reports that *ets-1* affects pigment cells in planarians (Fincher *et al*., 2018; He *et al*., 2017; Stubenhaus *et al*., 2016)(Fig S6B). After *ets-1*(RNAi), relative gene expression of pigment markers *pgbd-1* and *gst* were reduced by 12% and 28% relative to controls (Fig S6B).

We also examined transcript levels for several genes expressed broadly in *cathepsin^+^* cells—*forkhead box factor-1* (*foxf-1*) (Scimone et al., 2018), *cathepsin F* (*ctsf*), and *low density lipoprotein receptor related-3* (*ldlrr-3*) (Roberts-Galbraith *et al*., 2016). Compared to control, *ets-1*(RNAi) animals had significant reduction across all three *cathepsin^+^* cell markers (Fig S6C). The *cathepsin^+^* cluster (also known as the parenchymal cluster) also includes eight subclasses of cells or states that are uncharacterized, many of which express *ets-1* (Fincher *et al*., 2018; Plass *et al*., 2018)(Fig S6A). We repeated RT-qPCR using eight additional genes that mark individual *cathepsin^+^* subclusters. After *ets-1*(RNAi), expression of these genes was significantly reduced (*aqp1, dd_5690,* and *dd_9*), significantly increased (*dd_1831),* or unchanged (*TTPA, dd_7593, cathepsin L2*/*ctsl2*, and *protein tyrosine phosphate receptor type ptprt*) (Fig S6D-E). Our RT-qPCR data thus paint a complex picture of the roles for *ets-1* in *cathepsin^+^* cell types outside of pigment cells and glia, but we do not see universal downregulation of transcripts that fits neatly with the conclusion that *ets- 1*(RNAi) affects all cells in the *cathepsin^+^* cluster in the same way. We also re-analyzed previously published RNA-seq data on *ets-1*(RNAi) animals that utilized different RNAi and amputation paradigms (Dubey *et al*., 2022). We did not see consistent, significant downregulation of genes enriched in *cathepsin^+^* cell subtypes that would indicate a uniform loss of some or all *cathepsin*^+^ cell types (Fig S6F, Table S4).

We conclude that *ets-1* knockdown induces differences in gene expression across multiple *cathepsin^+^* subclusters that depends on the specific target gene, cell subpopulation, and amputation site chosen. While our data strongly implicate *ets-1* in maintenance and regeneration of planarian glia and confirm the role of *ets-1* in pigment cells, roles for *ets-1* in regulating other specific subclusters of *cathepsin*^+^ cell types will require further study. Importantly, glial cells are the only *cathepsin^+^* cell type present specifically in the nervous system, which allowed us to use *ets-1(RNAi)* as a first step for perturbing glia and examining neuronal organization and animal behavior.

### Reduction of *ets-1* affects CNS neuron gene expression and organization

Across metazoans, glial cells have been extensively implicated in neuronal development, maintenance, survival, and function. For example *Drosophila* glia regulate neuronal proliferation and, consequently, neuronal numbers (Coutinho-Budd *et al*., 2017; Ebens *et al*., 1993; Plazaola-Sasieta *et al*., 2019). Glial cells in *C. elegans* demarcate regions within the nervous system (*i.e.* nerve ring formation) before neuronal proliferation and consequently, also regulate neuron numbers (Rapti *et al*., 2017; Yoshimura *et al*., 2008). We showed that *ets-1*(RNAi) affects planarian glial gene expression and glial cell number. This discovery allowed us to investigate potential consequences of glial cell perturbation in planarians.

We first asked if perturbation of *ets-1* impacted gross morphology of the nervous system. We knocked down *ets-1* and examined expression of *ChAT* at 7dpa (Nishimura *et al*., 2010)(Fig 6A). We quantified brain area relative to body area and saw a significant decrease in brain size after *ets-1*(RNAi) (Fig 6B). Next, we examined whether neuronal gene expression was affected following head regeneration coincident with the loss of glia. We first utilized RT-qPCR to quantify relative gene expression of individual neuronal markers such as *ChAT*, *glutamic acid decarboxylase* (*gad*; Nishimura *et al*., 2008), *tyrosine hydroxylase* (*th*; Fraguas *et al*., 2012; Nishimura *et al*., 2007), *tryptophan hydroxylase* (*tph*; Nishimura *et al*., 2007), *neuropeptide precursor-3* (*npp-3*; Collins *et al*., 2010) and *secreted peptide prohormone -12* (*spp12;* Ong *et al*., 2016; Shimoyama *et al*., 2016) (Fig 6C). Interestingly, when we performed RT-qPCR to quantify *ChAT* gene expression, we did not see a significant decrease in transcript levels (Fig 6C). In contrast, *ets-1*(RNAi) animals had significantly increased mRNA levels for *gad*, and to a lesser extent *th* and *tph* (Fig 6C), all genes important for neurotransmitter biosynthesis. Neuropeptide-encoding mRNAs *npp-3* and *spp-12* were not affected by *ets-1*(RNAi) (Fig 6C).

**Figure 6.**
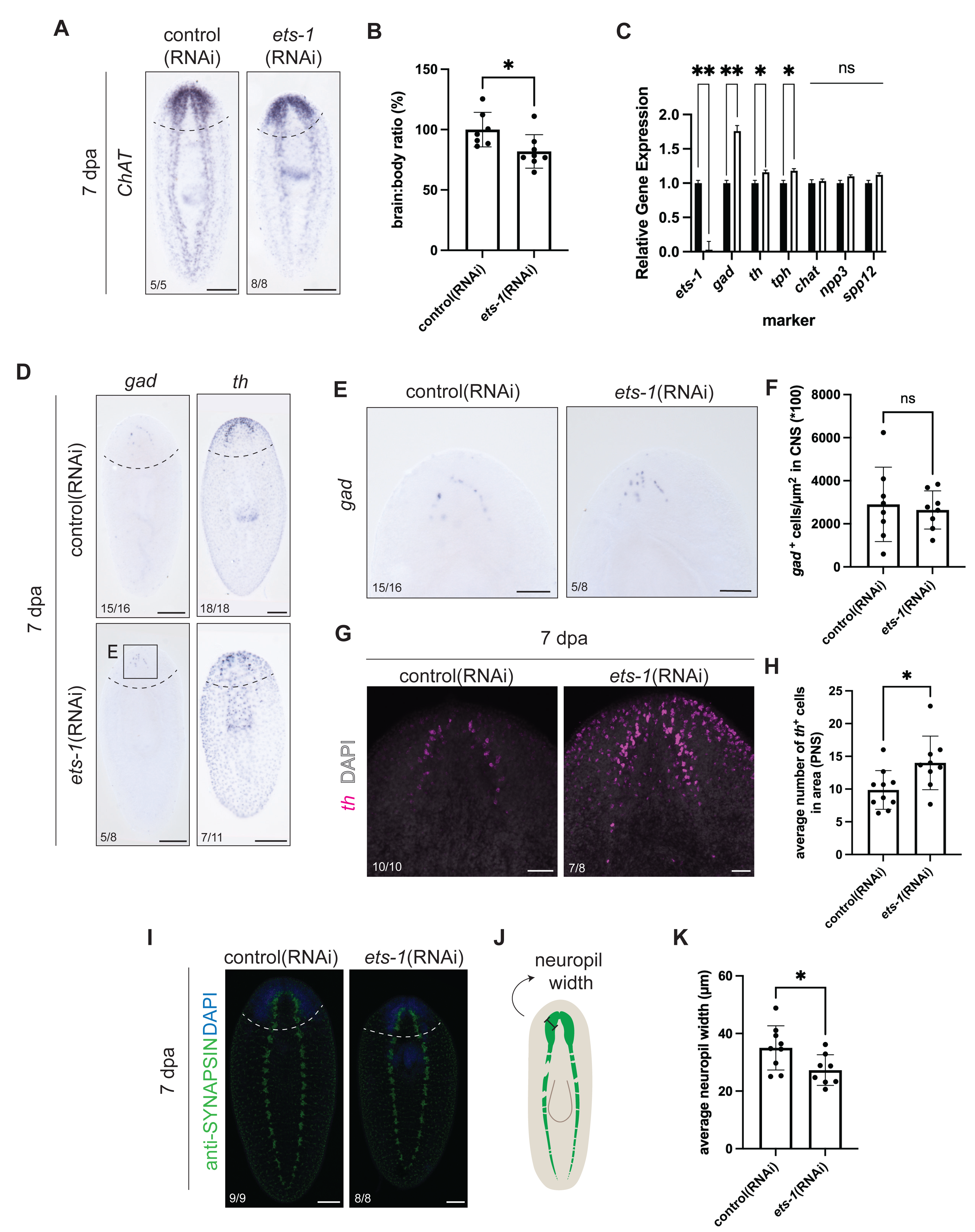
Knockdown of *ets-1* affects neuronal organization and gene expression. (A) Regenerated control and *ets-1*(RNAi) animals were subjected to ISH with riboprobes for *ChAT* (Nishimura *et al*., 2010). Dashed line depicts amputation site. (B) A brain-to- body ratio was calculated. *ets-1*(RNAi) animals exhibited significant reduction in brain area. *p≤0.05 (unpaired t-test with Welch’s correction). Error bars: SD. (C) RT-qPCR was used to detect levels of individual transcripts associated with neurotransmitter biosynthesis after RNAi. *p≤0.05, **p-value≤0.01, ns=not significant (unpaired t-tests with Welch’s correction). (D) ISH of regenerated control and *ets-1*(RNAi) animals with riboprobes for *glutamic acid decarboxylase* (*gad*) (Nishimura *et al*., 2008) or *tyrosine hydroxylase* (*th*) (Fraguas *et al*., 2012; Nishimura *et al*., 2007). Dashed line depicts amputation site. (E) Inset of *gad* expression in control and *ets-1*(RNAi) animals. 62.5% of *ets-1*(RNAi) animals had a disorganized *gad*^+^arch pattern. (F) Quantification of *gad*^+^ cells, normalized to body size, was compared. ns = not significant (unpaired t-test with Welch’s correction). Error bars: SD. (G) 7 dpa control and *ets-1*(RNAi) animals were subjected to FISH with riboprobes for *th* (magenta) and DAPI (grey). (H) *th*^+^cells in the PNS were counted in 100 µm^2^areas. *ets-1*(RNAi) animals had increased *th*^+^cell numbers in the PNS. *p≤0.05 (unpaired t-test with Welch’s correction). Error bars: SD. (I) Immunofluorescence with anti-Synapsin, which labels synapses in the neuropil and through the body, in control and *ets-1*(RNAi) animal (7 dpa; DAPI in blue). (J) Illustration of neuropil width. (K) Quantification of neuropil width. *p≤0.05 (unpaired t-test with Welch’s correction). Error bars: SD. Anterior up. Scale bars: (A, D) 200 µm (E, I) 100 µm (G) 50 µm.

We hypothesized that the increase in gene expression of *gad*, *th* and *tph* could be due to either 1) an increase in neuronal cell number or 2) an increase in mRNA levels without change in the number of cells. To differentiate between the two possibilities, we first performed ISH on *ets-1*(RNAi) animals after regeneration and examined expression of *gad* and *th* (Fig 6D). Interestingly, we noted that 62.5% of the *ets-1*(RNAi) animals had abnormal patterning of *gad*^+^ cells, where the linear, arched organization of *gad*^+^ cells seen in control animals was lost in *ets-1*(RNAi) animals (Fig 6E). To quantify *gad*^+^ and *th*^+^ cells in the regenerated heads, we repeated our RNAi paradigm and performed FISH on control and *ets-1*(RNAi) animals. We did not see a change in *gad*^+^ cell number in the regenerated head in *ets-1*(RNAi) worms compared to control (Fig 6F). Similarly, we did not see a change in *th*^+^ cell numbers in the CNS or pharynx (Fig 6D, G; S7A-B). However, we did see an increase in *th*^+^ cells in the PNS in *ets-1*(RNAi) animals in this experiment (Fig 6G-H). In addition, based on both colorimetric ISH and FISH data, we also consistently noted an increase in the intensity of *th* expression within each cell after *ets-1*(RNAi) (Fig 6D, G). Based on our data, we conclude that loss of *ets-1* impacts several neuronal mRNAs but perturbs cell number for only some subsets of neurons (PNS *TH+* cells).

Taken together, our data show that the reduction of *ets-1* results in: decreased brain size, altered organization of specific cell types (*gad*), anomalous neuronal gene expression, and change in cell number for some neuron types (PNS *TH*+). We did not see phenotypes that would indicate widespread neuronal cell death or failure to specify neurons during regeneration after reduction in glial number. Our data together indicate that inhibition of *ets-1*, possibly due to loss of glial cells, broadly but subtly impacts planarian neuronal gene expression and CNS organization.

### Loss of *ets-1* does not lead to broad loss of neural connectivity

Glial cells in other animal species play important roles in synapse organization and function (Eroglu & Barres, 2010; Lee & Chung, 2019; Neniskyte & Gross, 2017). To determine if *ets-1* knockdown affected synapse density or organization in planarians, we stained animals 7dpa with *anti-*Synapsin (Fig 6I) (Ross *et al*., 2015). We quantified medial brain gap (the space between the two brain lobes), average VNC gap length, average gap sizes, and average width of the neuropil, a synapse- and process-rich structure in the interior of the planarian brain (Fig 6J; S7C-F). *ets-1*(RNAi) worms did have a significant decrease in neuropil width compared to control (Fig 6K). However, we saw no additional effects on organization or staining intensity in the synapses of *ets- 1*(RNAi) worms compared to control (Fig S7D-F). Reduction in neuropil size could explain our previous observation of a small brain size after *ets-1*(RNAi) without decrease in neuronal cell number (Fig 6B). In addition, we also note that the neuropil is the location of most glial cells in the planarian CNS, suggesting that loss of glial cells could impact local organization of neuronal processes or synapses in this region.

Glia also play important roles in axon guidance, axon fasciculation, and axonal targeting in other species (Hidalgo *et al*., 1995; Mason & Sretavan, 1997; Poeck *et al*., 2001). In planarians, individual axon trajectories can be most clearly seen in the visual system. Furthermore, we also noted that glial marker *estrella* can be seen near the eyespots (Fig 3A) (Roberts-Galbraith *et al*., 2016). To determine if glial cells play roles in photoreceptor axon trajectory in planarians, we performed *ets-1*(RNAi) to reduce glial cell number and stained with *anti-*Arrestin (Sakai *et al*., 2000) (Fig S7G). We quantified gaps in the optic chiasm and numbers of stray bundles or axons near the photoreceptors, optic chiasm, and neuropil (Fig S7H-J). We did not see any significant differences in photoreceptor axon structure in *ets-1*(RNAi) animals. We concluded that *ets-1* does not affect overall axonal morphology in the photoreceptor system, although roles for axon guidance in other neurons will need to be investigated once better tools are developed.

### Perturbation of *ets-1* results in locomotion defects

We reasoned that reduction in glial cell number might also affect neuronal *function*. We rationalized that impaired neuronal function could be reflected in planarian behavior, including response to light. Planarians normally exhibit negative phototactic behavior, preferring to move toward areas of low light (Fig 7A; S8; Video 1; based on assays in (Paskin *et al*., 2014; Zewde *et al*., 2018)). *ets-1*(RNAi) animals consistently showed reduced ability to move into a dark space compared to control animals (Fig 7A- B; S8; Videos 1A-B). By 5 minutes, more than 50% of *ets-1*(RNAi) animals remained on the light side of the dish and by 10 minutes, 41.18% of *ets-1*(RNAi) animals failed to reach the dark side (Fig 7B; Video 1B). In comparison, 75% of control worms reached the dark side within 3 minutes, and by 6 minutes, over 96% of control worms had reached the dark side (Fig 7B; S8; Video 1A). Interestingly, we noticed that *ets-1*(RNAi) worms often exhibited uncoordinated movements that included head lifts and inch- worming, a slow locomotion gait based on muscle contraction instead of cilia-mediated movement (Video 1B). When we quantified inch-worming behavior, 61.76% of *ets- 1*(RNAi) animals exhibited inch-worming behavior as they initiated movement compared to control animals (9.38% displayed inch-worming behavior) (Fig 6C). Even in the absence of a light gradient, 23.4% of *ets-1*(RNAi) animals exhibited inch-worming behavior at movement onset compared to 4.35 % of control animals (Fig 6D). We conclude that *ets-1*(RNAi) impacts both the quality and outcome of planarian movement, reflecting possible glial roles in robust neuronal function.

**Figure 7.**
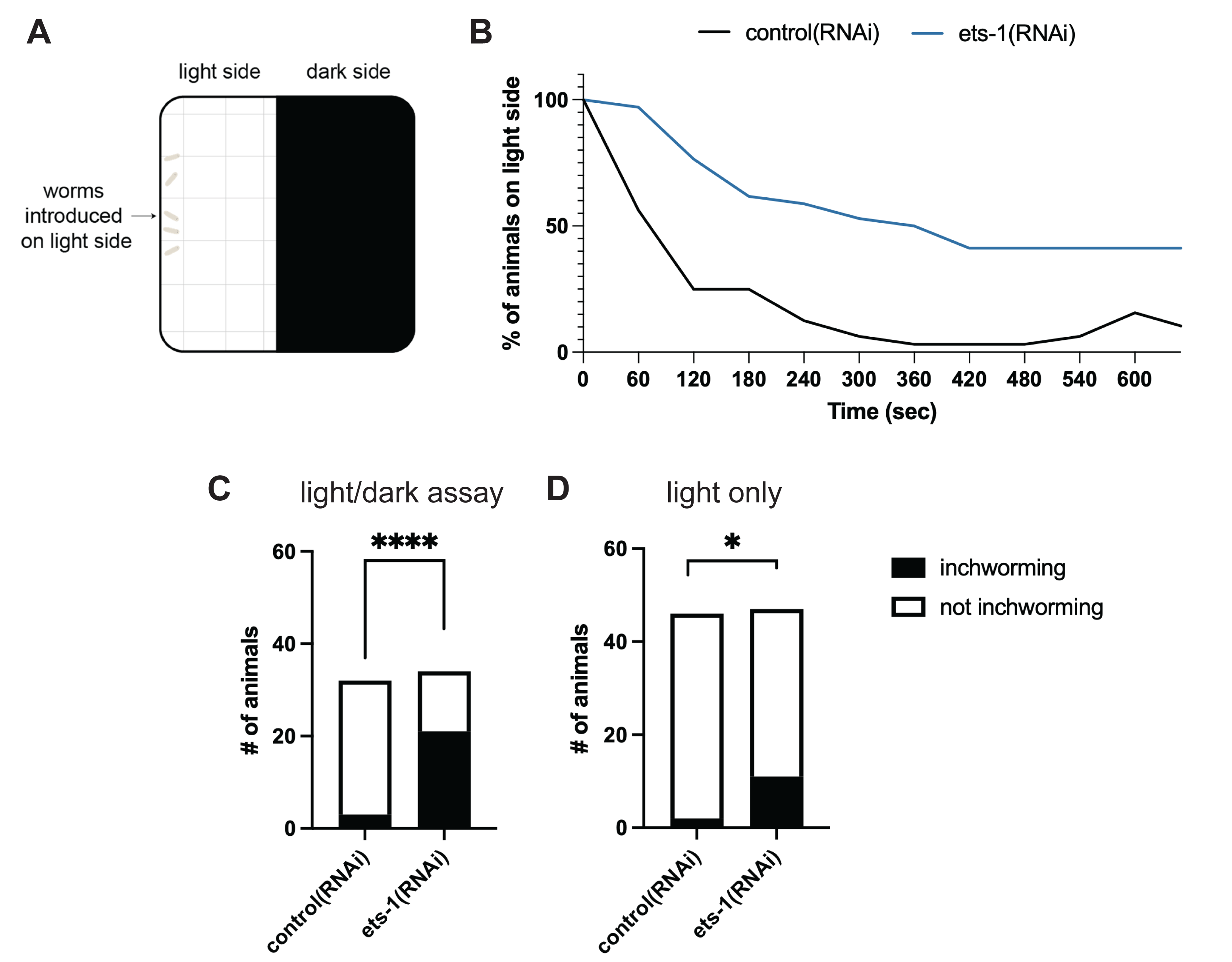
*ets-1* knockdown leads to changes in planarian behavior. (A) Illustration of light/dark assay (Paskin *et al*., 2014; Zewde *et al*., 2018). (B) A graph showing percentage of intact animals that remained on the light side over 600 seconds (n=10-12 animals per replicate; 3 replicates). (C) Quantification of animals that exhibit inch-worming behavior in the context of a light/dark assay. More *ets-1*(RNAi) animals exhibited inch-worming behavior during or prior to movement to the dark side of the dish. ****p≤0.0001 (Fisher’s exact test). (D) In the absence of a light gradient, *ets- 1*(RNAi) still led to a significantly higher incidence of inch-worming behavior compared to control. *p≤0.05 (Fisher’s exact test).

## Discussion

Planarians regenerate their nervous systems quickly and with high fidelity, making them an attractive model system for studying glia during successful nervous system repair. In this work, we established timelines for development and regeneration of glial cells, demonstrating that glia arise after neurons and depend on neurons or pigment cup cells for placement in the region of the eye. We further show that transcription factor *ets-1* plays conserved roles in gliogenesis and glial maintenance in *Schmidtea mediterranea*. Our work further capitalized on our findings with *ets-1* to explore potential roles of glial cells in planarians for the first time. Our results indicate that *ets-1*(RNAi) causes altered gene expression in neuronal cell types, reduction in neuropil size, and perturbed fluidity of animal movement. It is important to note that *ets- 1* affects multiple *cathepsin*^+^ cell types in planarians. Further work will be required to dissect the roles of Ets-1 in diverse cell types and to more specifically ablate glia to confirm and expand possible role(s) for glia in regulating neural physiology and behavior. Moreover, it will be interesting to address how *ets-1* specifies glial fate and to explore the identity and function of Ets-1 targets in planarian glial cells.

### The role of *ets-1* in gliogenesis is conserved in planarians

Members of the *Ets* family have conserved roles in driving gliogenesis across metazoans. ETS transcription factors are important for both CNS and PNS gliogenesis (Hagedorn *et al*., 2000). *Drosophila* Pointed, an ETS transcription factor expressed in glial cells, is required for differentiation of longitudinal and midline glial cells (Klaes et al., 1994; Klämbt, 1993). *Ets-1* is expressed in human cortex astrocytes and plays roles in astrocyte differentiation, proliferation, and regulation of genes involved in astrocyte signaling (Amouyel *et al*., 1988; Fleischman *et al*., 1995). ETS family transcription factors are also crucial for survival and proper regeneration of mammalian Schwann cells and zebrafish bridging glia after injury (Arthur-Farraj *et al*., 2012; Klatt Shaw *et al*., 2021; Nagarajan *et al*., 2002; Parkinson *et al*., 2002). Similarly, we found that planarian *ets-1* is also required for glial regeneration and maintenance. *ets-1* knockdown reduces glial numbers, with differential effects on individual glial markers.

Planarian *ets-1* also affects pigment cells (He *et al*., 2017) and we showed that it regulates gene expression in other cell types in the planarian body, most of which are not well characterized. Interestingly, planarian glial cells clustered more closely with pigment cells than neurons in single-cell transcriptomic analyses (Fincher *et al*., 2018). This finding argues against common progenitors for neurons and glia in planarians and indicates that glia might share a common progenitor with other phagocytic, *cathepsin^+^* cell types (*e.g.* pigment cells). Alternatively, *ets-1* could regulate cell state in a wide variety of phagocytic, *cathepsin^+^* cell types that are not lineage-related but that cluster together due to similarities in gene expression that correspond to functional rather than lineage relationships.

### Relationship between planarian glia and neurons

During CNS development in vertebrates and *Drosophila*, neurogenesis often precedes gliogenesis as stem cells produce neurons before switching programs to make glia (Campbell & Götz, 2002; Miller & Gauthier, 2007; Molofsky & Deneen, 2015). This sequence of events is shared across species with varying time scales. Here, we uncovered a similar “birth order” in planarian regeneration and development, in which neurons arise first and glia arise later. While vertebrates and *Drosophila* possess a neurogenic-to-gliogenic switch during neural development, planarian neurons and glial cells are thought to arise from distinct lineages of pluripotent stem cells (Crews, 2019; Fincher *et al*., 2018; Miller & Gauthier, 2007; Plass *et al*., 2018; Roberts-Galbraith *et al*., 2016; Wang *et al*., 2016). Interestingly, though pluripotent stem cells give rise to all cells in the adult planarian body, this is the first observation to our knowledge of a “birth order” of cell types that contribute to a common tissue in planarian. Our observations raise new questions regarding why and how planarian stem cells produce glia and neurons with different timing after injury.

### Potential roles for glial cells in the planarian nervous system

Glial cells fulfill diverse roles across animals including regulating neuronal cell numbers and migration, aiding in axon guidance and growth, assisting in neuron differentiation, regulating synapse formation and pruning, regulating ion homeostasis, providing metabolic support, participating in sensory systems, and promoting and/or impeding response to injury (Allen & Lyons, 2018; Falk & Götz, 2017; Jäkel & Dimou, 2017; Oikonomou & Shaham, 2011; Ortega & Olivares-Bañuelos, 2020; Shaham, 2015; Yildirim *et al*., 2019).

Our studies with Ets-1 allowed us to investigate potential roles for planarian glia for the first time. Our data suggest that planarian glia promote brain organization. We observed disorganized *gad*^+^ cells after *ets-1* knockdown, suggesting that glial cells might play a role in proper pattering for specific neuron types or in establishing boundaries between neurons during head regeneration. In addition, we observed reduction of neuropil width after *ets-1*(RNAi). The neuropil is densely packed with processes and synapses of neurons and is also the location of the highest concentration of glial cells, suggesting a possibility of planarian glia assisting in synapse maturity, organization, or function. Further tools to study synaptic activity (*i.e.*, calcium signaling) will be essential for investigation of glial effects on neurotransmission and synapse function. Conversely, our data do not support essential or broad roles for planarian glial cells in regulating neuronal numbers or neuronal survival.

Glial expression of *glutamine synthetase* and *excitatory amino acid transporter* leads us to hypothesize that planarian glial cells regulate neurotransmitter uptake and recycling (Roberts-Galbraith *et al*., 2016; Wang *et al*., 2016). We also note that when *ets-1* was knocked down and glial cells are reduced, there was an increase in *gad* expression but no change in neuronal number in the CNS. One possible explanation for this observation is that *gad*^+^ neurons may alter gene expression in response to persistence of extracellular neurotransmitters in the absence of recycling by glial cells (Araque *et al*., 1999; Oliet *et al*., 2001).

Additionally, planarian glia are likely to exhibit phagocytic properties, based on their classification as *cathepsin*^+^cells (Fincher *et al*., 2018) and functional analysis (Scimone *et al*., 2018). Vertebrate glia (notably astrocytes, microglia and Schwann cells), *Drosophila* glia, and *C. elegans* glial cells phagocytose apoptotic neurons and neurite debris during development and following neuronal injury (Aldskogius & Kozlova, 1998; Jessen & Mirsky, 2016; Jung & Chung, 2018; Logan & Freeman, 2007; Sulston *et al*., 1983). However, new tool development will be necessary to assess the purpose of phagocytosis by planarian glia as well as additional roles for glia in modulating neurons.

### Planarian glial cells and behavior

Our data indicate that loss of glial cells is associated with changes in planarian behavior, based on our finding that *ets-1*(RNAi) impacts the quality and speed of movement in planarians. This locomotion defect was more pronounced when *ets- 1*(RNAi) were presented with a more complex visual stimulus and when animals initiated movement. We know little about whether planarian glia may impact movement locally at the neuromuscular junction (NMJ), or whether glia impact integration of sensory information, decision making, and initiation of movement through neurons in the CNS. In vertebrates, the NMJ is composed of a presynaptic motor neuron terminal, a post-synaptic muscle cell, and perisynaptic glial cells (typically Schwann cells) that cooperate for motor output (Reddy *et al*., 2003). Further ultrastructural work could reveal whether planarian glia reside near synapses between neurons and/or between neurons and other cell types.

More work will also be required to determine the basis of movement defects in *ets-1*(RNAi) animals. One possibility is that dysregulation of neurotransmitter abundance in *ets-1*(RNAi) animals impacts negative phototaxis. In particular, *gad^+^* GABAergic neurons and *TH+* dopaminergic neurons have been implicated in movement in planarians (Nishimura *et al*., 2007; Nishimura *et al*., 2008). Further work could explore whether neurotransmitter levels are impacted in the absence of glia and whether exogenous neurotransmitters or antagonists could rescue behavioral defects in *ets-1*(RNAi) animals.

### Planarian glial cells can help us understand the evolutionary origin and diversification of glia

Discovery and characterization of glial cells in emerging model organisms present unique challenges. Molecular markers have often been used to identify cell types and have become a prevalent tool for cell identification in the ‘omics era. However no glial genes are universally expressed among all glial types or across all species.

Thus, identification of glial cells in new species remains challenging. Previous works have shown the importance of ETS transcription factor in both glial development and regeneration in vertebrates (Hagedorn *et al*., 2000; Kiyota *et al*., 2007; Parkinson *et al*., 2002). Our discovery that *ets-1*, a transcription factor with conserved roles in gliogenesis, also plays an important role in glial cell regeneration in planarians suggests that *ets-1* or downstream targets of *ets-1* could be potential candidates for molecular markers that could be used to identify novel glial cells in other animal models.

Finally, a thorough characterization of planarian glia will fill gaps in our understanding of glia in an underexplored, highly regenerative phylum. Future work may reveal fascinating new aspects of glial biology, and provide insight into glial evolution, development, and regeneration. Our work with planarian glia provides a valuable point of comparison and contrast with glial cells in other organisms, particularly in the areas of glial function and glial response to injury.

## Acknowledgements

We are grateful for the assistance from University of Georgia Biology Microscopy Core and Dr. K.M. Kandasamy. We thank Kendall Clay, Jennifer Jenkins, and Taylor Medlock-Lanier for thoughtful feedback on this work and manuscript. anti-Synapsin antibody was obtained from the Developmental Studies Hybridoma Bank, created by the NICHD of the NIH and maintained at The University of Iowa. We are grateful to Dr. Alejandro Sánchez Alvarado and Shane Merryman (Stowers Institute) for planarians provided during our laboratory start-up. We are grateful to the farmers at White Oak Pastures (Bluffton, GA) for raising healthy cattle that we used for our beef liver. This research was supported by the Alfred P. Sloan Foundation (RRG), the McKnight Foundation (RRG), the University of Georgia Department of Cellular Biology, and the Center for Cancer Research, National Cancer Institute, National Institutes of Health Intramural Research Program project number ZIABC0120009 (ED). The content of this publication does not necessarily reflect the views or policies of the Department of Health and Human Services, nor does mention of trade names, commercial products or organizations imply endorsement by the US Government.

---

**Video 1. *ets-1*(RNAi) animals have changes in negative phototaxis behavior.**

(A) Control animals before head amputation in light/dark assay, 20x playback speed. (B) *ets-1*(RNAi) animals before head amputation in light/dark assay, 20x playback speed. Video only shows the light side of the dishes.

**Video 2. ets-1(RNAi) animals exhibit locomotion defects.**

(A) Control animals before head amputation with no stimulus, 20x playback speed. (B) *ets-1*(RNAi) animals before head amputation with no stimulus, 20x playback speed.

**Supplemental Table 1.** List of cloning primers.

**Supplemental Table 2:** List of protein sequences for phylogeny tree.

**Supplemental Table 3.** List of RT-qPCR primers.

**Supplemental Table 4.** Metadata of qPCR and RNA-seq of *ets-1*(RNAi) across multiple gene expression across in *cathepsin*^+^cells *(Dubey et al., 2022; Fincher et al.*, 2018; Plass *et al*., 2018).

## Methods

### Animal Husbandry

A clonal line of diploid, asexual *Schmidtea mediterranea* (CIW4 strain) was maintained at 18-22°C in the dark. Animals were kept in Ziploc® (9 cup) reusable containers in 1X Montjuïc salts as previously described (Cebrià & Newmark, 2005). Organic, pureed calf or beef liver (White Oak Pastures, Georgia) was used to feed animals once per week. Animals were starved for a minimum of 1 week prior to use in experiments. For sexual *Schmidtea* husbandry and embryo staging, outbred cohorts of sexually reproducing planarians descended from animals collected in Sardinia by Dr. Maria Pala (1999) and Drs. Longhua Guo and Alejandro Sánchez Alvarado (2015) were reared in 1X Montjuïc salts (Cebrià & Newmark, 2005) at 20°C in constant darkness. Sexually mature animals were housed at low density and fed homogenized beef liver (White Oak Pastures, Georgia) twice per week. To promote fertility, breeding adult sexual *Schmidtea* stocks were replaced every 3 months with young adults (6-8 weeks post-hatching) or adult regenerates (6-8 weeks post-amputation). Egg capsules were collected daily, soaked in 10% bleach for 3 minutes, rinsed 4-6 times, and stored in 1X Montjuïc water in a 20°C incubator.

### Identification of genes and cloning

Planarian homologs of genes of interest were identified from PlanMine 3.0 (Rozanski *et al*., 2019) based on homology and from scRNA-seq data (Fincher *et al*., 2018). Primers shown in Table S1 were designed using Primer3 (Rozen & Skaletsky, 1999) to PCR-amplify 500-750 bp segments of genes of interest from asexual *S. mediterranea* cDNA synthesized from iScript kit (Bio-Rad)(Collins *et al*., 2010). Each PCR product was ligated into Eam1105I-digested pJC53.2 vector for use in RNAi and ISH experiments using standard molecular biology protocols (Collins *et al*., 2010).

### Protein alignment and Phylogenetic Analysis

Protein alignment and phylogenetic analysis were performed as described in (Jenkins & Roberts-Galbraith, 2022). Briefly, the longest open reading frame for each sequence was identified using web-based translation tool Expert Protein Analysis System [ExPASy, https://www.expasy.org/; (Gasteiger, 2003)] or NCBI Open Reading Frame Finder (https://www.ncbi.nlm.nih.gov/orffinder/). Protein sequences of ETS-1 from other species were aligned to reference sequences (Table S2). Phylogeny was performed using www.phylogeny.fr (Dereeper *et al*., 2008) using MUSCLE for sequence alignment with the “A la Carte” option (Edgar, 2004) and PhyML for phylogenetic tree construction (Guindon *et al*., 2010).

### RNAi experiments

For RNAi experiments, animals (10-12 worms; 3-4 mm in size) were kept in 60 mm Petri dishes, washed after feeding and supplemented with 1:1000 gentamicin sulfate (50 mg/mL stock, [Gemini Bio-Products]) throughout the experiment. dsRNA was synthesized using standard molecular biology techniques (Collins *et al*., 2010). dsRNA matching *Aequorea victoria green fluorescent protein* (*GFP*) was used for negative control feedings. Animals were fed 5-10 µg dsRNA mixed with 25-30 µL of 3:1 beef liver:1X Montjüic salts mixture and 2 µL McCormick^®^ green food dye. Feedings were completed every 3 days for a total of 5-6 feedings. For regeneration experiments, animals were amputated pre-pharyngeally seven days after the last feeding and fixed according to different downstream protocols. For long-term RNAi experiments, 30 animals per RNAi condition were fed 5 µg or 10 µg dsRNA every 3 days for a period of 60 days. The number of animals alive was quantified as detailed in Fig S3A. Survival curves were plotted using GraphPad Prism9.

### *in situ* hybridization (ISH) and immunofluorescence (IF)

Animals utilized for ISH were treated with 7.5% N-acetyl-L-cysteine (NAC) in Phosphate Buffered Saline (PBS: 1.37 M NaCl, 27 mM KCl, 100 mM Na2HPO4,20 mM KH2PO4, pH 7.4), and fixed in 4% formaldehyde (in PBSTx: PBS + 0.3% Triton-X 100). Riboprobes for ISH on asexual planarians were generated using PCR-amplified products from pJC53.2 vectors with T7 primers (orientations provided in Table S1). Antisense probes were synthesized with digoxigenin-11-UTP (Roche) or fluorescein-12-UTP (Roche) using standard molecular protocols (Collins *et al*., 2010). ISH experiments were performed on asexual animals as previously described (King & Newmark, 2013). Some samples were processed in an Insitu Pro (Intavis) hybridization robot. Probes were detected using anti-digoxigenin antibodies with Fab fragments (Sigma-Aldrich) that were conjugated with an alkaline phosphatase (Roche) for colorimetric ISH or horse-radish peroxidase (Roche) for fluorescent ISH (FISH). For colorimetric ISH, 5-Bromo-4-chloro- 3-indolyl phosphate (BCIP, [Roche]) and nitro blue tetrazolium chloride (NBT, [Roche]) in alkaline phosphatase (AP) buffer were used for signal development. Animals were mounted in 80% glycerol. Samples were imaged with an Axiocam 506 color camera mounted on Zeiss Axio Zoom V.16 microscope using ZEN 2.3 pro software.

For FISH, probes were detected using anti-digoxigenin and anti-fluorescein antibodies with Fab fragments conjugated to horseradish peroxidase (Roche). 1:500 fluor-tyramide (TAMRA or FAM), 1:1000 4-IPBA, and 0.003% H2O2 in Tyramide Signal Amplification (TSA) buffer (2 M NaCl; 0.1 M Boric acid, pH 8.5) was used for signal development for 45 minutes. Double FISH samples were incubated in Sodium Azide solution (100 mM sodium azide [Fisher Scientific] in PBS + 0.3% TritonX-100 (PBSTx)) to inactive peroxidase activity for 45 minutes before performing secondary signal development.

FISH samples were mounted in VectaShield^®^ Antifade Mounting Medium and imaged using a Zeiss LSM 880 Confocal microscope with an upright AXIO Imager Z2 and ZEN Black 2.3 SPI software. Whole animals were imaged using a Plan-Neofluar 10X/0.3 objective and smaller fields were imaged with Plan-Apochromat 20X/0.8 objective (no immersion). For each channel, the cut-off between signal and background was determined using the fluorescent intensity range indicator. Images shown are representative of all images taken in an experiment.

Embryo staging was performed according to guidelines set forward previously (Davies *et al*., 2017). The collection date was considered 1 day post-egg capsule deposition (dped). Early spherical Stage 6 (S6) embryos were fixed at 6-7 dped; elongating mid-stage 6 (mid-S6) embryos were fixed at 8 dped; Stage 7 (S7) embryos were fixed at 10 dped; Stage 7.5 (S7.5) embryos were fixed at 12 dped; and Stage 8 (S8) embryos were fixed at 14-15 dped using the protocol described in (Davies *et al*., 2017) for colorimetric ISH with the following modifications: Embryos were removed from egg capsules immediately prior to fixation. S6 and S6.5 embryos were fixed for 6 hours to overnight in 4% paraformaldehyde in PBSTx (PBS+0.5% Triton X-100) at room temperature. S7, S7.5 and S8 embryos were treated with 5% NAC in PBS (S7: 2 minutes; S8: 4 minutes), immediately followed by fixation in 4% paraformaldehyde in PBSTx (PBS+0.5% Triton X- s1. 100) for 45 min (S8) or 2 hours (S7, S7.5) at room temperature. Colorimetric WISH images were acquired on a Zeiss Axio Zoom V16 equipped with a Axiocam 305 color camera. For image processing, a polygonal lasso tool was used to extract images of embryos from the original TIFFs; these were transferred to a white background along with the scale bar. Brightness and contrast were adjusted on colorimetric images to facilitate visualization of the colorimetric ISH signal.

Briefly, for immunofluorescence experiments, asexual planarians were treated in 2% HCl, fixed for 15 minutes in 4% formaldehyde in PBSTx (PBS+0.3% TritonX-100), and then bleached in 6% H2O2 in PBSTx overnight. Animals were blocked in 1% Bovine serum Albumin (BSA) in PBSTx overnight at 4°C. The primary antibodies used were anti-Synapsin (1:100, 3C11 concentrate; Developmental Studies Hybridoma Bank) and anti-Arrestin (1:1000, cat #016-arrestin-01, LagenLabs). Secondary antibodies were used at a dilution of 1:500-1:1000 (goat anti-mouse Alexa Fluor 488 and goat-anti-rabbit Alexa Fluor 488, respectively [Invitogen]) (Ross *et al*., 2015; Sakai *et al*., 2000).

Samples were mounted in VectaShield^®^ Antifade Mounting Medium and imaged using a Zeiss LSM 880 Confocal microscope with ZEN Black software. Whole animals were imaged using 10X/0.3 objective, specific regions were imaged with 20X/0.8 objective (no immersion).

### Analysis of RNA sequencing data

Analysis for early time point regeneration of asexual and sexual planarian glial expression was performed from previously published RNA-sequencing data (Davies *et al*., 2017; Roberts-Galbraith *et al*., 2016) and plotted using GraphPad Prism9. Re- analysis of *ets-1*(RNAi) gene expression across different *cathepsin*^+^ subclusters was performed using previously published RNA-sequencing data (Dubey *et al*., 2022) and expression profiles provided in previous transcriptome data analysis (Fincher *et al.,* 2018; Plass *et al*., 2018).

### Real-time quantitative PCR (RT-qPCR)

Total RNA was isolated from RNAi-treated animals using Trizol (Invitrogen) as per the manufacturer’s protocol. 1 µg of RNA was reverse transcribed into cDNA using an iScript cDNA Synthesis Kit (BioRad) as per manufacturer’s protocol. RT-qPCR was completed using Applied Biosystems QuantStudio3 Real-Time PCR system using GoTaq qPCR Master Mix with SYBR Green (Promega). All primers used for RT-qPCR are shown in Table S3. All measurements were performed in biological and technical triplicates; RNAi and RNA purification were performed from 3 individual Petri dishes (10- 12 worms each, biological triplicates) for each RNAi condition, and then we completed each RT-qPCR in three identical reactions per sample/primer pair (technical triplicates). Overall transcript normalization was accomplished using *beta-tubulin* mRNA within each sample. Statistical analyses were performed using GraphPad Prism9; details of each statistical test are provided in the figure legends.

### Quantification of ventral nerve cord gaps, brain to body ratio, cell numbers and neural structures

#### Quantification of VNC gaps (*estrella* staining)

For quantification of ventral nerve cord(VNC) gaps in *estrella* expression, FIJI (Schindelin *et al*., 2012) was utilized to measure overall VNC length per animal by summing the length of the left and right VNC. The length of each true gap (defined by absence of *estrella* expression in VNC) was summed per animal and averaged. Then, a ratio of gap length to total VNC length per animal was determined.

#### Quantification of cell numbers

For quantification of *estrella*^+^ cells in *ets-1*(RNAi) head regenerates, cells were counted manually in the new blastema for each animal and averaged. For quantification of *estrella*^+^ cells in the trunk pieces in *ets-1*(RNAi), *estrella*^+^ cells were counted for 5 separate pre- and post-pharyngeal 200 µm^2^areas per animal and averaged, as previously described (Stelman *et al*., 2021). For quantification of *estrella*^+^*, calamari*^+^*, estrella*^+^/*calamari*^+^, *estrella*^+^/*calamari*^-^, *estrella*^-^/*calamari*^+^ cells in *ets-1*(RNAi); *estrella*^+^ and *pkd2L-1*/*pkd1L-2*^+^ cells in *soxb1-2(*RNAi), FISH images were quantified in Bitplane IMARIS 9.9 (Oxford Instruments) using spots or colocalization modules. Cells were counted in the new head blastema or specified regions for each animal within 250 µm^2^or 200 µm^2^areas, respectively. For quantification of *gad*^+^ cells, cell numbers were manually counted in head blastemas from colorimetric ISH images. For quantification of *th*^+^ cells, 3D renditions of FISH images using FIJI were used to count cell numbers manually or automatically by 3D objects counter plugin and then corrected manually. For quantification of *estrella*^+^ cells around eye, z-stack images were analyzed using ImarisViewer 9.9.1 (Oxford Instruments) and then used to manually quantify the presence or absence of *estrella*^+^ cells in the region of the typical eye (left and right) for each animal per condition.

#### Quantification of brain size

Brain to body ratios were determined by tracing the *ChAT*^+^ expression in the brain using FIJI and comparing brain area to body area, as previously described (Roberts-Galbraith *et al*., 2016; Schindelin *et al*., 2012).

#### Quantifications of neural structures

For quantification of neural structures in anti-Synapsin and anti-Arrestin immunofluorescence images, identities of samples were masked and randomized. Traits were quantified from maximum intensity projection images. For anti-Arrestin, samples were quantified for number of stray bundles or axons near the photoreceptors, stray bundles or axons in/near optic chiasma, stray bundles or axons near the neuropil, gaps in axon trajectory. For anti-Synapsin, samples were quantified for neuropil width (brightest parts and including fainter edges), brain gaps, and VNC gaps using FIJI. Brain width and brain gap criteria were measured in triplicate and averaged for each individual animal. For quantification of VNC gaps, the number of gaps on the left and right side was averaged and then divided by the total length of the animal (in mm) to get the average gap length per mm. To calculate the average gap length, measurements of each gap per individual sample were taken and then averaged per animal. The samples were then unmasked, and measurements were averaged for each RNAi condition. All statistical analyses were conducted using GraphPad Prism9; details of each statistical test are shown in the figure legends.

### Behavioral Assays

Prior to light/dark behavioral assays, RNAi animals were placed in VWR Square Petri Dishes (Electron Microscopy Sciences,13x13 mm) after the last dsRNA feeding to acclimate to the dishes for two days. For the light/dark assay, new Petri dishes were half covered with black electrical tape (Duck Brand^®^) on the lid and the corresponding side of the petri dish. Then 10-12 animals per RNAi condition were introduced at the edge or corner of the light side of the petri dish and recorded on an iPhone® 11 (Apple) from a height of approximately 17 centimeters from the top of the dish for approximately 10 minutes. The behavioral test was repeated in biological triplicate for each experimental condition. To quantify, the number of animals remaining on the light side was recorded at 60 second intervals. 0 seconds indicates when animals were introduced to the dish. Then, the percentage of animals (over three biological replicates) for each RNAi condition residing in the light side was calculated for each time point and plotted using GraphPad Prism9. For publication, videos were sped to 20X using Adobe Photoshop.

---

**Supplemental Figure 1.**
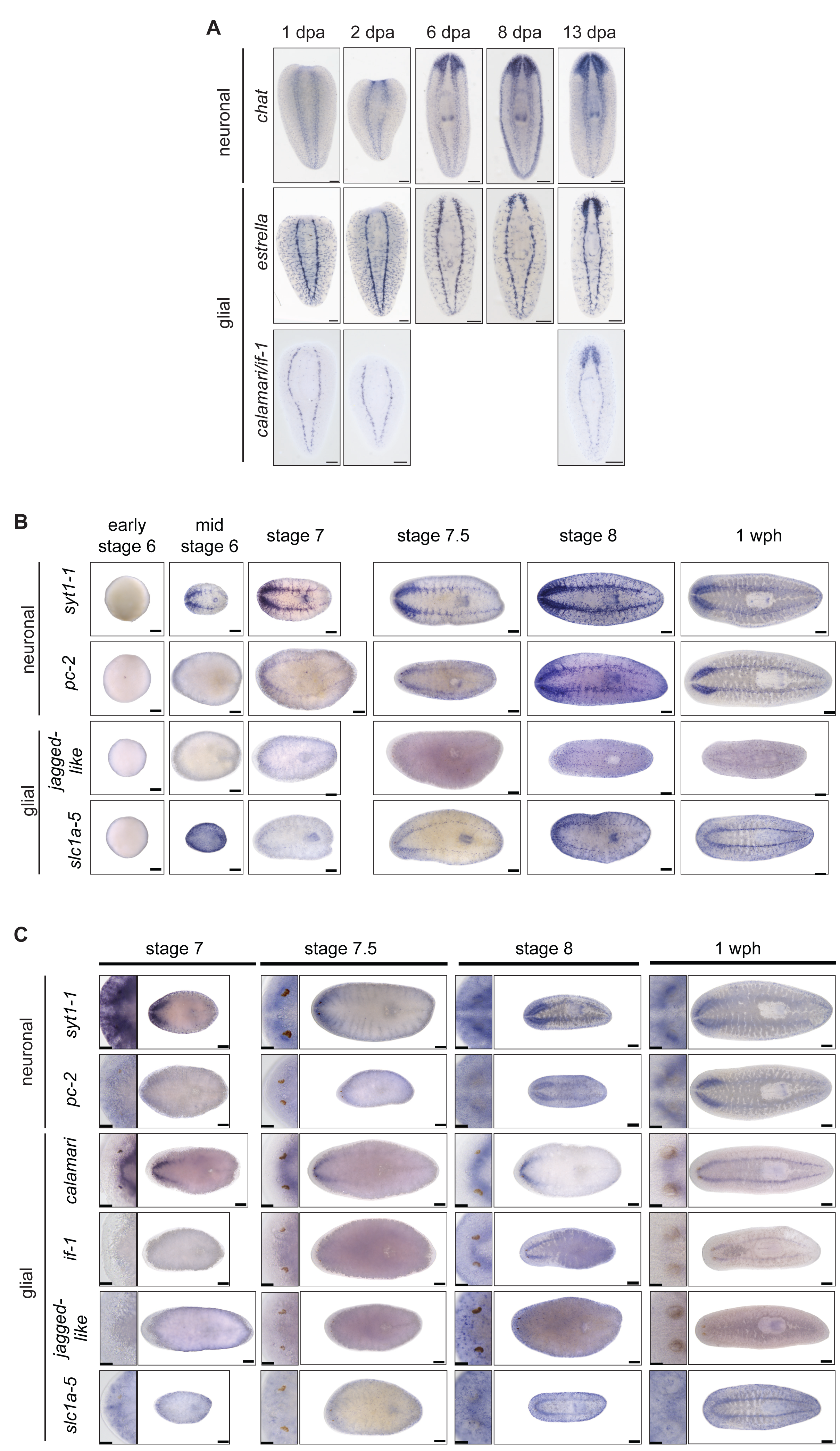
Neuronal and glial gene expression during head regeneration. (A) ISH regeneration timeline of neurons (*ChAT*) and glial markers (*estrella*, pooled *if- 1*/*calamari*) in asexual planarians at 1-, 2-, 6-, 8-, and 13-days after head amputation. Ventral view, anterior up, from same time series as Fig 1A. (B) ISH of neuronal markers (*pc-2, syt1-1*) and glial markers (*slc1a-5/EAAT, jagged-like*) on planarian embryos (early S6, mid-S6, S7, S7.5, S8) and juveniles (1 week post-hatching). Anterior to left, ventral views. (C) ISH of neuronal markers *(pc-2, syt1-1)* and glial markers (*if-1*, *calamari*, *jagged- like*, *slc1a-5/eaat*) at indicated developmental stages. Dorsal views. Insets show expression near the eyes. Scale bars: whole animal images 200 µm; eyes 50 µm.

**Supplemental Figure 2.**
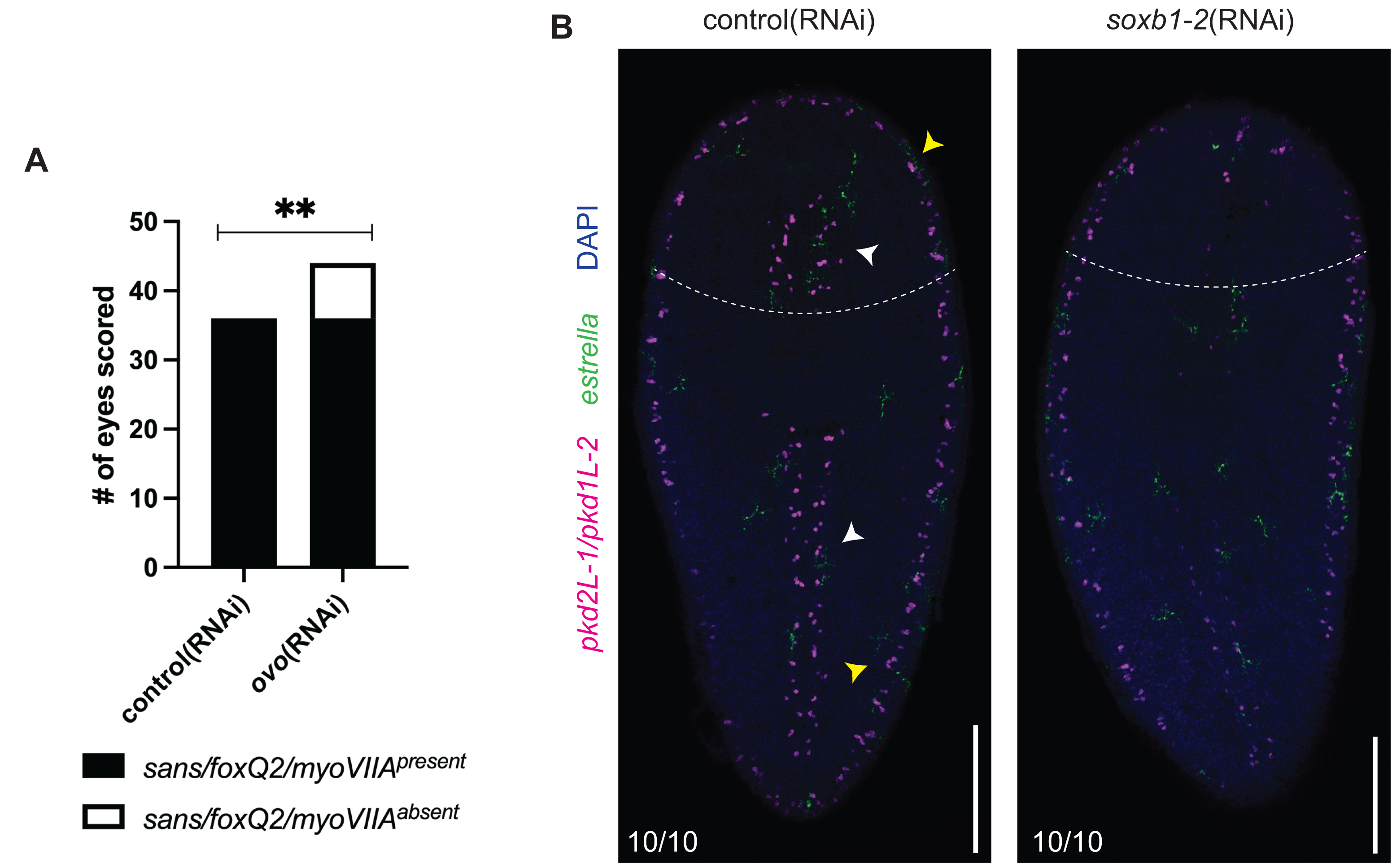
Loss of sensory *pkd*^+^ neurons does not affect glial cells in non-regenerating tissue. (A) Quantification of photoreceptor neuron markers (pooled *sans*/*foxQ2*/*myoVIIA*) present (black) or absent (white) in control and *ovo*(RNAi) eyes. Left and right eyes are considered separately. **p-value≤0.01 (Fisher’s exact test). (B) 7 dpa control and *soxB1-2*(RNAi) animals subjected to FISH detecting pooled *pkd2L-1*/*pkd1L-2* (magenta) and *estrella* (green), and DAPI (blue). Arrowheads indicate where dorsal *estrella*^+^ cells were observed: medial stripe (white) and lateral margin (yellow). Dashed lines indicate amputation site. Whole body animal from Figure 3. Dorsal views, anterior up. Scale bar: 200 µm.

**Supplemental Figure 3.**
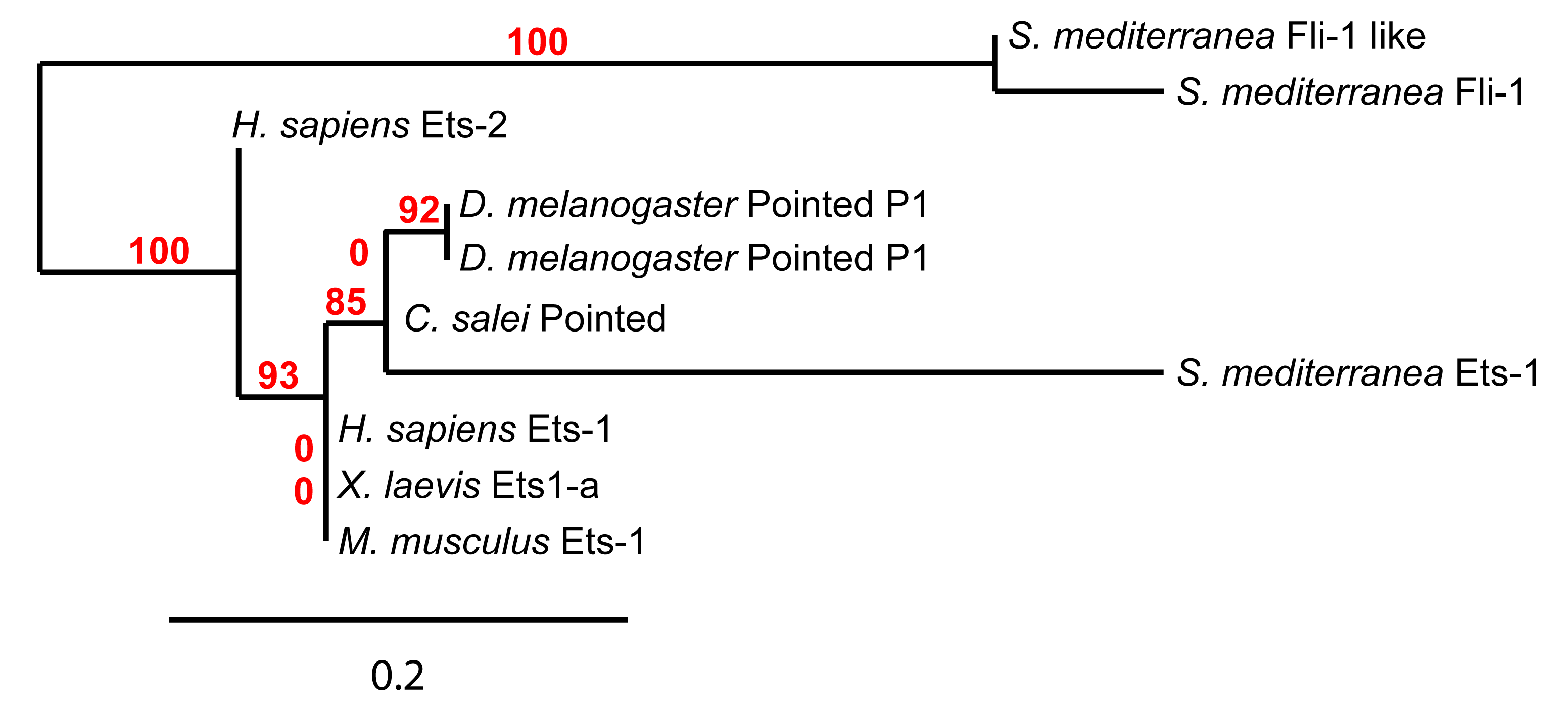
Ets-1 protein sequence is highly conserved. Phylogenetic tree of Ets-1 protein sequence of 6 species based on longest open reading frame. Analysis shows the relationship between planarian Ets-1 and Ets-1 protein sequences in other species. The outgroup is planarian Fli-1 and Fli-1-like proteins, two proteins that possess an ETS domain, but have no established role in glial cells. Red text denotes the percentage of support for each node.

**Supplemental Figure 4.**
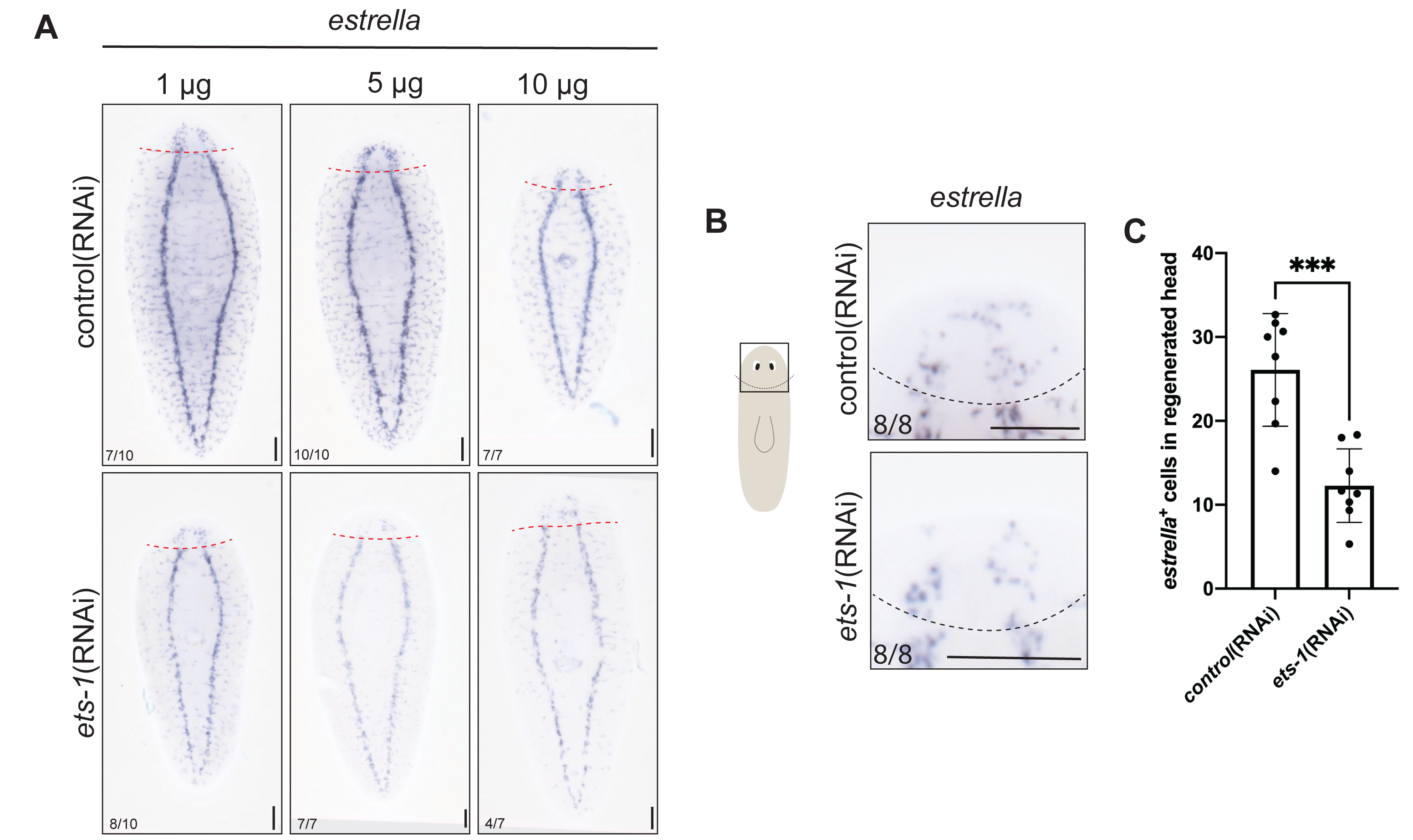
Ets-1 knockdown phenotype on glial markers varies depending on dosage. (A) Animals were fed 1 µg, 5 µg or 10 µg *in vitro*-transcribed dsRNA targeting *ets-1* for 5 feeds 3 days apart, amputated, fixed at 7 dpa, and subjected to ISH with *estrella*. 5 µg and 10 µg regimens yielded more robust reduction of *estrella*^+^cells in regenerated heads in *ets-1*(RNAi) animals. (B) *ets-1*(RNAi) resulted in fewer *estrella*^+^ cells present in 7 dpa regenerated head compared to control. (C) Quantification of *estrella^+^* cells in regenerated head blastemas in control and *ets-1*(RNAi) animals. ***p-value≤0.001 (unpaired t-tests with Welch’s correction). Dashed lines indicate amputation site. Scale bar: 200 µm.

**Supplemental Figure 5.**
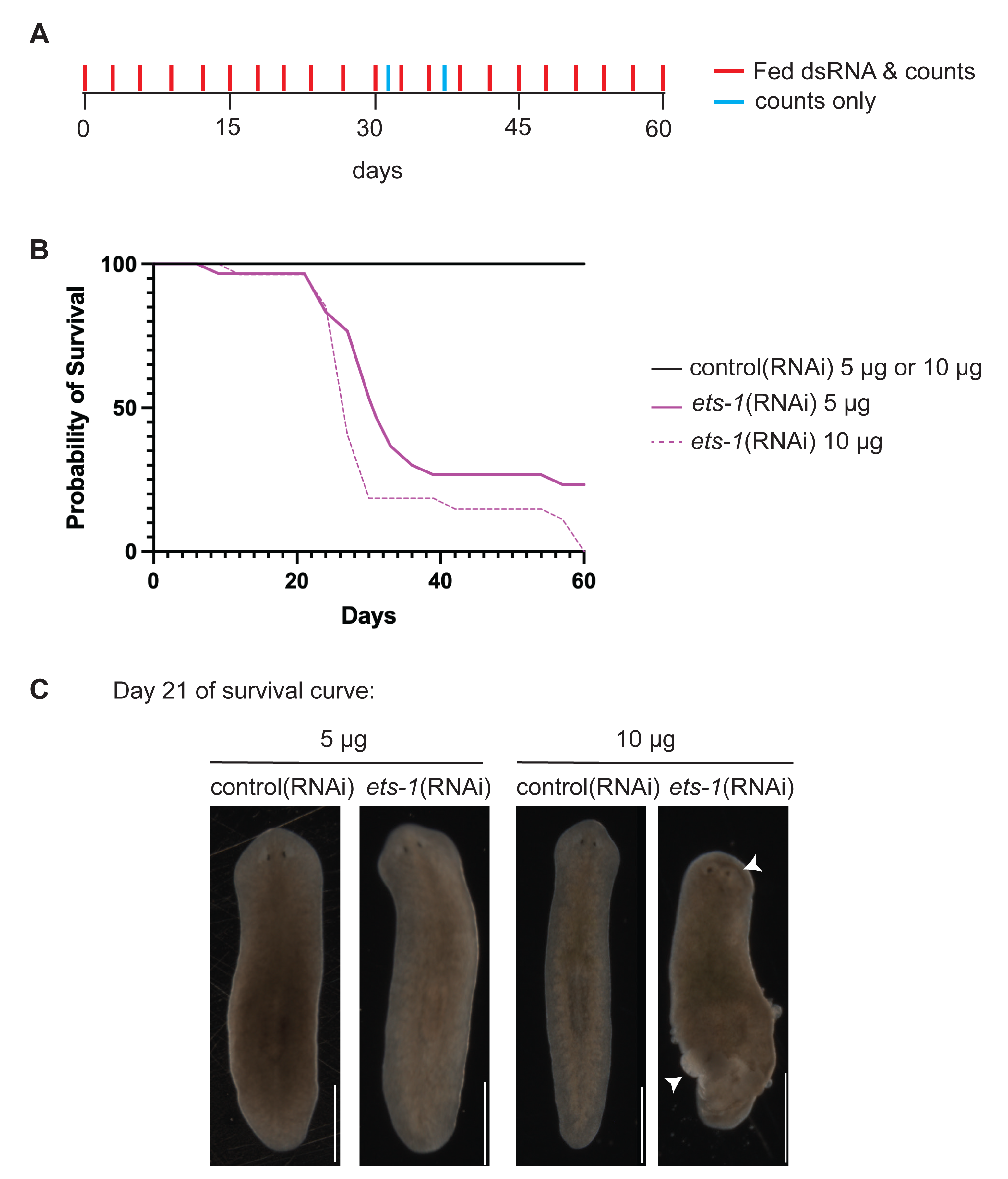
E*t*s*-1* knockdown leads to eventual animal death. (A-B) Animal cohorts were subjected to a 60-day dsRNA-feeding regimen, with feeds every 3 days. Red: Days with feeding and animal counts are marked in red. Blue: days with animal counts only. Survival curve depicting the relative percentage of surviving animals at different dsRNA feeding doses after long term RNAi; N=30 animals each. (C) Live images at day 22 (post 8^th^feeding) of control and *ets-1*(RNAi) animals fed dsRNA at dosages of 5 µg and 10 µg. *ets-1*(RNAi) animals at 10 µg exhibited lesions (white arrowheads) in the tail and around eyes that led to eventual lysis at later time points.

**Supplemental Figure 6.**
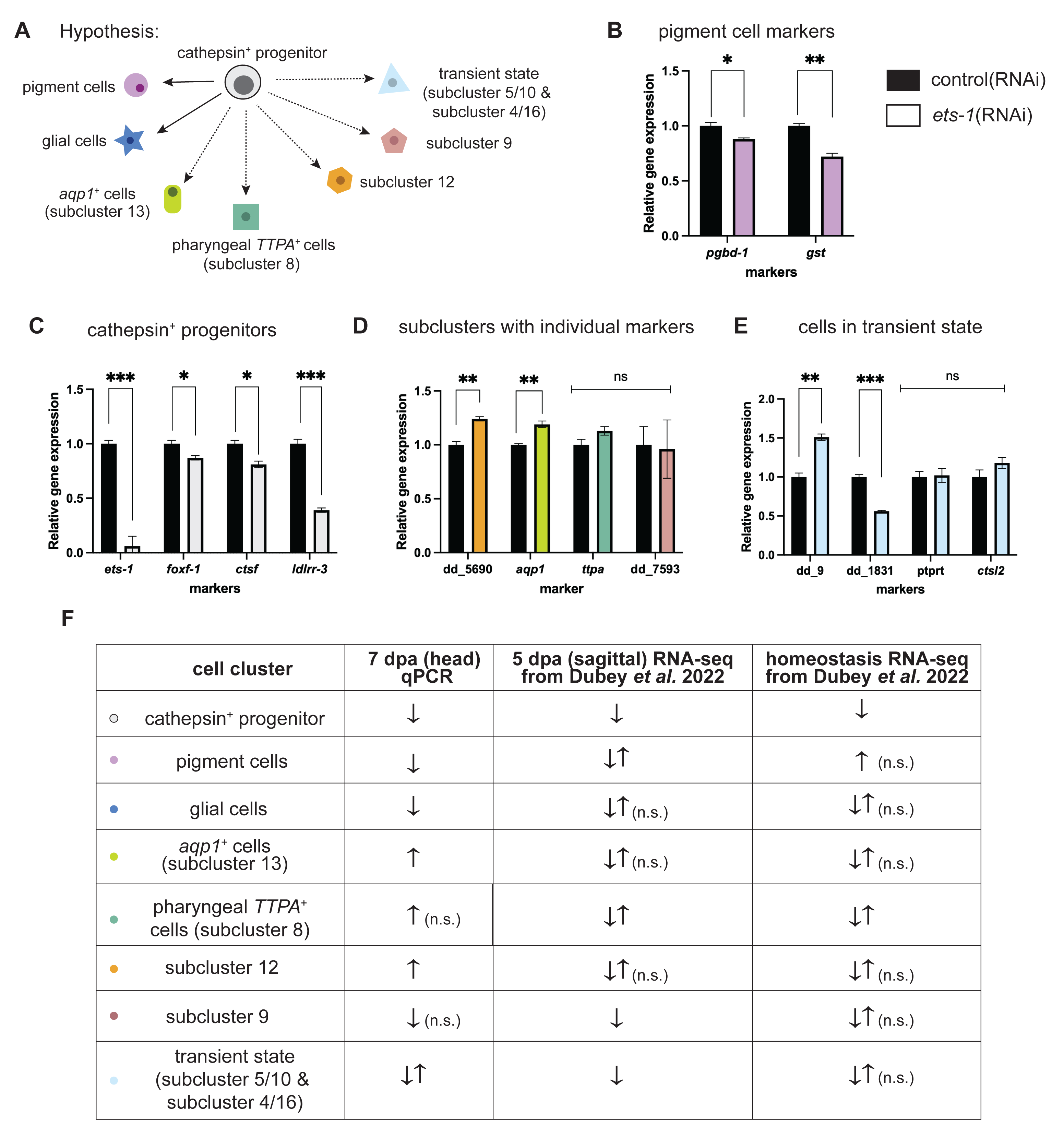
e*t*s*-1* affects gene expression in individual *cathepsin*^+^ cell types in distinct ways. (A) Graphical illustration of *cathepsin*^+^ cell subclusters identified in single cell transcriptomic atlases (Fincher *et al*., 2018; Plass *et al*., 2018). Clusters are color-coded as follows: grey = cathepsin^+^ progenitor cells, purple = pigment cells, dark blue = glial cells, light green = *aqp1^+^*cells, green = *TTPA^+^*cells, orange = subcluster 12, pink = subcluster 9, light blue = subclusters 5/10 and subcluster 4/16. Lineage relationships are drawn as previously proposed. (B-E) RT-qPCR was used to detect levels of *ets-1* and other markers of *cathepsin*^+^subclusters after RNAi with color coding as in A. Details of each marker are provided in Table S3. *p≤0.05, **p-value≤0.01, ***p- value≤0.001, ****p-value≤0.0001, ns=not significant (unpaired t-tests with Welch’s correction). Error bars: SEM. (F) Summary table of trends seen after *ets-1* knockdown in each cathepsin^+^ cell type from qPCR data and published RNA-seq data (Dubey *et al*., 2022). Details of each gene analyzed under each subcluster are provided in Table S4.

**Supplemental Figure 7.**
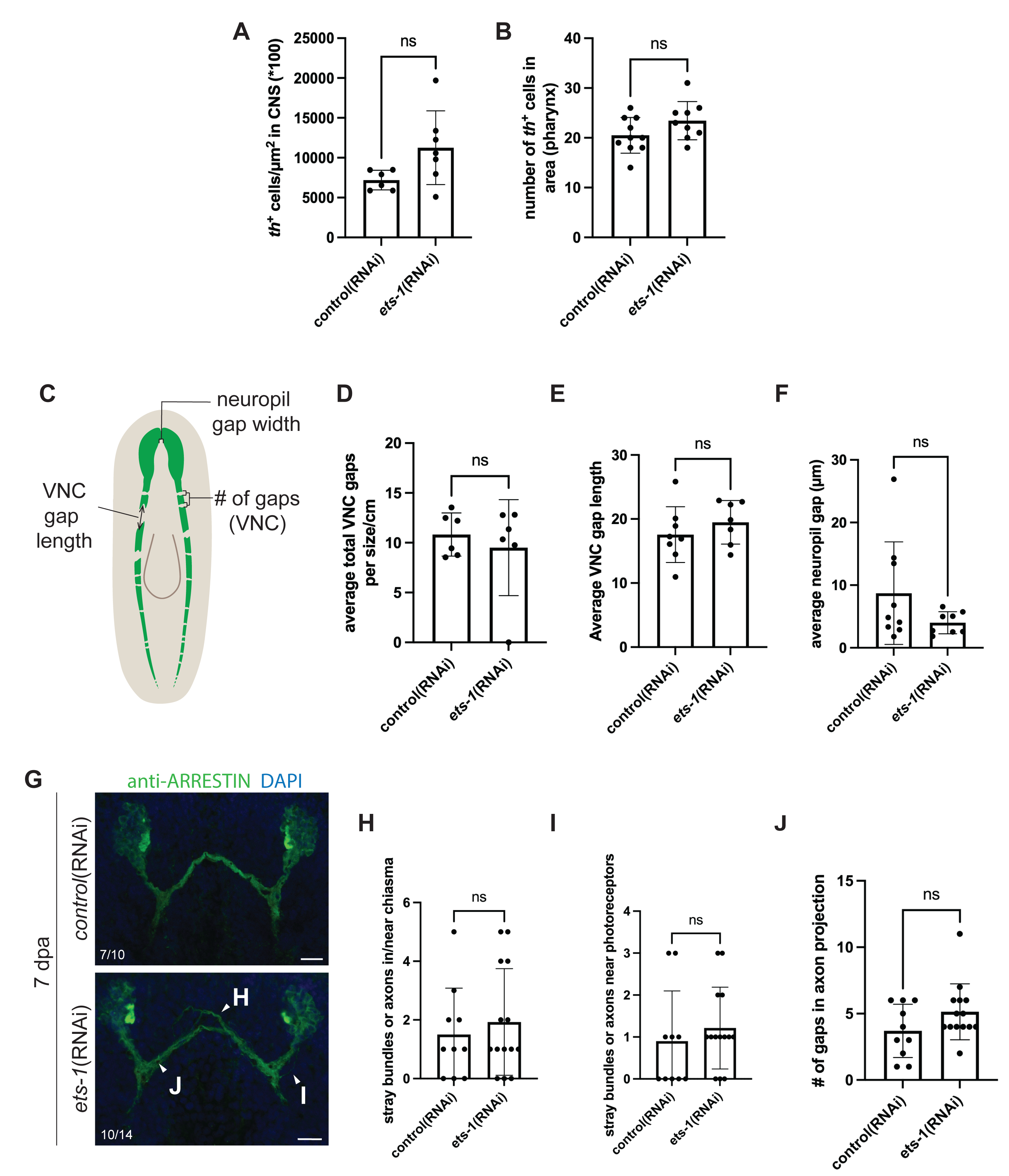
e*t*s*-1* knockdown does not affect neural architecture. (A) Quantification of *th*^+^ cells in the brain normalized to body size in *ets-1*(RNAi) animals and compared to control animals. *ets-1* knockdown resulted in a non-significant increase in *th*^+^ cell numbers in the CNS. ns=not significant (unpaired t-test with Welch’s correction). Error bars: SD. (B) Quantification of *th*^+^ cells in the pharynx in 100 µm^2^areas in control and *ets-1*(RNAi) animals. *ets-1* knockdown did not change *th*^+^ cell in the pharyngeal nervous system. ns=not significant (unpaired t-test with Welch’s correction). Error bars: SD. (C) Graphical illustration of criteria that were quantified from anti- Synapsin immunofluorescence images: gap length, number of gaps, and neuropil gap length. (D-F) Quantification of total gaps in VNC per cm, average gap size in VNC (µm), and average gap length between neuropil (µm) for control and *ets-1*(RNAi) animals. (G) Immunofluorescence of anti-Arrestin (green) labels photoreceptor axon projections in control and *ets-1*(RNAi) at 7 dpa. DAPI in blue. (Bottom) White arrows highlight (H) fraying at the optic chiasm, (I) secondary axon projections to the eye and (J) gaps (or “holes”) in the axon projection bundle. (H-J) Quantification of stray bundles or axons in or near the optic chiasm, near photoreceptors (left and right), and gaps in axon projections in control and *ets-1*(RNAi) animals. Each point is an individual animal. ns=not significant (unpaired t-test with Welch’s correction for unequal variances). Error bars: SD. Scale bar: 20 µm.

**Supplemental Figure 8.**
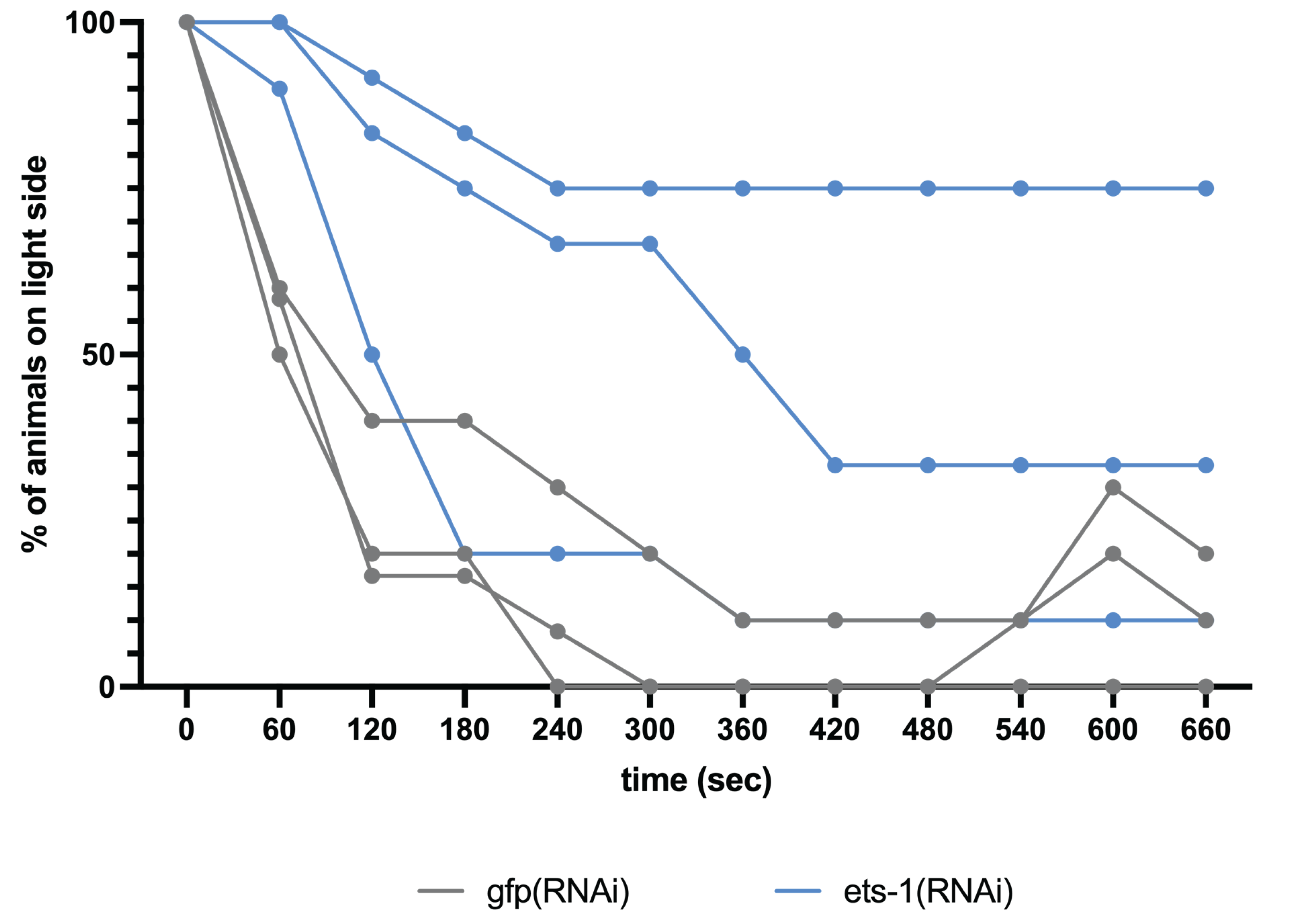
Individual replicates of *ets-1*(RNAi) animals exhibited changes in phototaxis behavior. Graph showing percentage of animals from individual replicates of control and *ets- 1*(RNAi) animals that remained on the light side during filming across 600 seconds. Three replicates were completed (n=10-12 per replicate).

